# TFEB activation ameliorates listeriolysin O-mediated subversion of lysosomal and LC3 function during *Listeria monocytogenes* infection

**DOI:** 10.1101/2025.11.30.691302

**Authors:** Aarti Pant, Sneha Bhatt, Parul Sen, Deepam Bhattacharya, Kuldeep Singh Rautela, Mamta Negi, Ravi Manjithaya

**Affiliations:** Autophagy Laboratory, Molecular Biology and Genetics Unit, Jawaharlal Nehru Centre for Advanced Scientific Research, Jakkur, Bangalore 560064, India

**Keywords:** *Listeria monocytogenes*, Xenophagy, Listeriolysin O, TFEB, Lysosomes, LC3, LIR and Autophagy

## Abstract

Xenophagy is a crucial innate subcellular defense mechanism, functioning as a selective autophagy pathway that captures intracellular pathogens within xenophagosomes for subsequent lysosomal degradation. However, many pathogens have evolved strategies to subvert this process, ensuring their intracellular replication and survival within the host cell. Despite its significance, the precise mechanisms by which intracellularly motile xenophagy cargo like *Listeria monocytogenes* (*Lm*) circumvent this selective autophagy pathway are not yet completely understood. In this study, we identify a previously unrecognized function of the secreted virulence factor listeriolysin (LLO) in orchestrating a multi-layered disruption of host autophagy-lysosomal machinery. While LLO is classically known for its role in mediating vacuolar escape, our findings reveal that it also contains a functional LIR motif which enables LLO to directly bind LC3 in the cytosol, rendering it unavailable for xenophagosome formation. Concurrently, LLO impairs lysosomal function by reducing its hydrolytic capacity and suppresses nuclear translocation of TFEB, a key transcription factor that governs autophagy and lysosome biogenesis, further compromising host degradative pathways. Using various *Lm* mutant strains, we show that actin-based motility is dispensable for xenophagy evasion. Instead, the intracellular persistence of *Lm* primarily stems from two factors: LLO-mediated LC3 binding, which diverts LC3 from its xenophagic functions, and the inhibition of TFEB nuclear translocation. Importantly, pharmacological activation of TFEB or genetic disruption of the LLO-LC3 interaction restores xenophagy and substantially reduces bacterial load. Collectively, our findings position LLO as a multifaceted virulence determinant that employs previously uncharacterized mechanisms to subvert host cell-autonomous defense. This study deepens mechanistic insight into the pathogenesis of *Lm* and highlights LC3 and TFEB as promising therapeutic targets for host-directed therapies against intracellular infections.

**Highlights:**

▪ *Listeria monocytogenes* (*Lm*) evades xenophagy independently of its host-actin polymerization ability
▪ Listeriolysin O (LLO) impairs lysosomal degradative potential and suppresses TFEB nuclear translocation
▪ LLO binds LC3 through its LC3-interacting region (LIR) motif, effectively limiting LC3 availability for xenophagosome formation
▪ Restoring LC3 and TFEB function enables intracellular clearance of *Lm*

**Graphical abstract:** The graphical abstract illustrates how vacuolar (Δ*hlyA*), and cytosolic (WT and Δ*actA*) *Listeria monocytogenes* differentially engage with host-degradative pathways. Listeriolysin O (LLO), a critical virulence factor, is highlighted as the linchpin determining whether these bacteria get targeted for degradation or successfully evade the host response. In Δ*hlyA* strain infected cells, where LLO is absent, (1) the bacteria remain confined within the phagosome. (2 - 4) The phagosomal compartment then fuse with lysosome, to form phagolysosome that is both acidified and hydrolytically active, enabling effective bacterial degradation while sustaining autophagy flux. (5) This process coincides with nuclear translocation of the master transcription regulator, TFEB, which enhances lysosomal biogenesis and function, boosting bacterial clearance. On the other hand, both WT and Δ*actA* strains produce LLO, (1, 2) which allows them to escape from the phagosome into the host cytosol. (3) WT strain induces polymerization of host cell actin, leading to the formation of actin clouds that mature into actin tails, facilitating bacterial mobility and cell to cell spread. (4) The Δ*actA* strain, despite lacking the ability to interact with host actin, still replicates in the cytosol. LLO plays a multifaceted role in evading host defenses. (5 - 6) It limits LC3 availability for nascent xenophagosome formation by binding to LC3, reducing bacterial capture by the xenophagy machinery. (7 - 8) Furthermore, even when a few bacteria are successfully sequestered within phagosome or xenophagosome, LLO-mediated lysosomal dysfunction impair their maturation into degradative phagolysosome and xenolysosome. (9) Compounding this, LLO suppresses nuclear translocation of TFEB, thereby dampening transcriptional activation of autophagy-lysosomal pathways. Collectively, these mechanisms compromise the host degradative capacity, allowing cytosolic bacteria to replicate and evade clearance. Several unanswered questions remain regarding the structural and mechanistic aspects of the interaction of LLO with LC3. It is still unclear whether LC3 conjugation occurs on single or double membrane vesicles, and whether these vesicular structures are fully sealed. Additionally, it is not yet determined if LLO binds LC3 specifically at sites of omegasome or phagophore formation. The aggregation behavior of LLO also raises intriguing possibilities: could these aggregates play a protective role in shielding LLO from its degradation? Furthermore, it remains to be clarified whether LLO exists freely in the cytosol or is associated with specific membrane-bound compartments. Finally, the precise mechanism by which LLO interferes with TFEB nuclear translocation is another critical open question requiring further investigation.

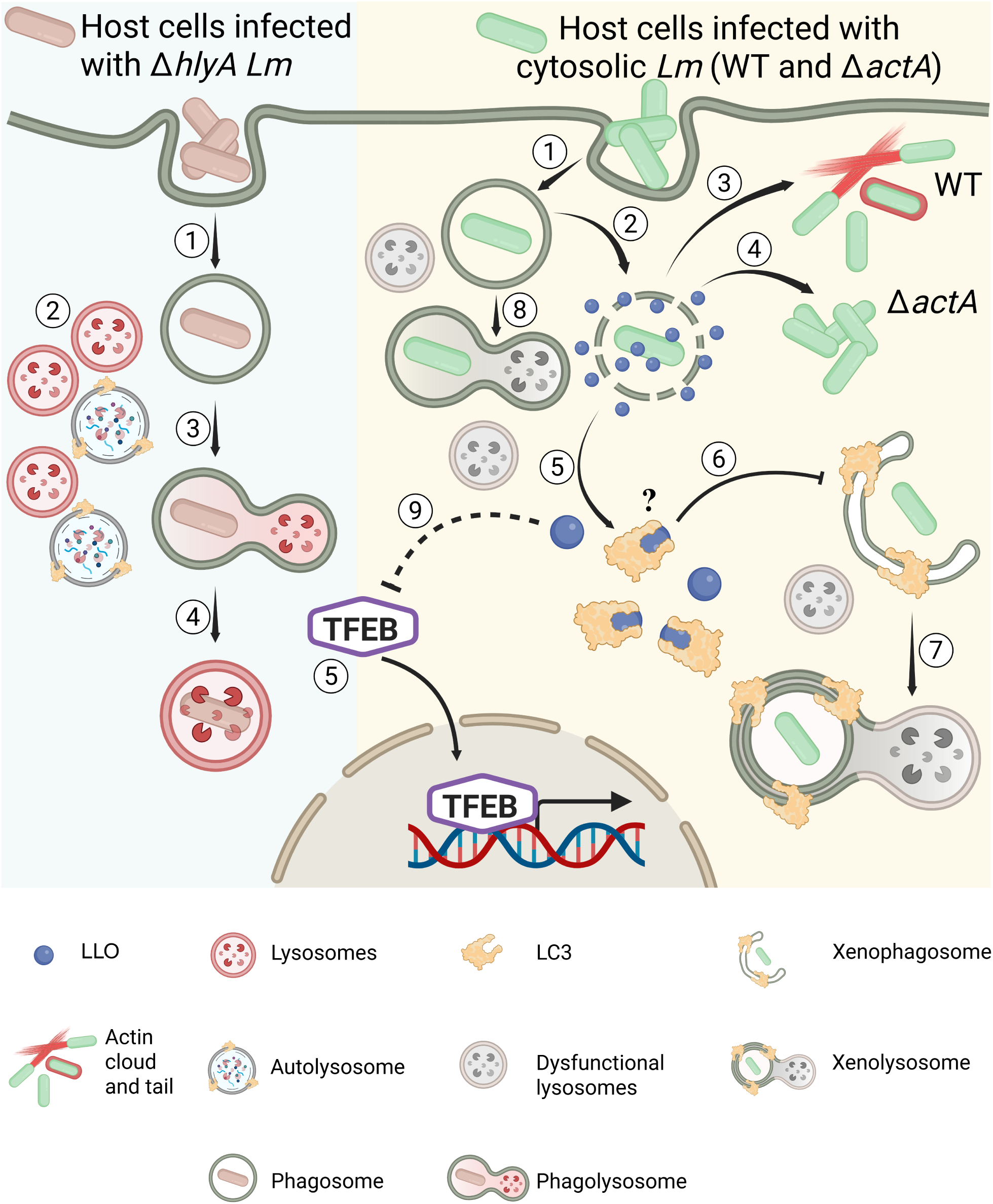

## Introduction

Our cells maintain homeostasis by intricate quality control mechanisms, chief among them is the autophagy-lysosomal pathway ^1^. Macroautophagy, hereafter referred to as autophagy, a fundamental catabolic process responsible for dismantling and recycling damaged organelles and macromolecules ^2,3^. A specialized form of autophagy, xenophagy, plays a critical role in our innate immune response by specifically targeting and degrading intracellular pathogens within autophagy-lysosomal compartment ^3–13^. Multiple studies suggest that xenophagy is regulated by a master transcriptional regulator, Transcription Factor EB (TFEB), which governs the expression of genes involved in lysosomal biogenesis, autophagy initiation, and autophagosome-lysosome fusion ^14–16^. In certain intracellular infections, TFEB has been shown to translocate to the nucleus, thereby inducing a transcriptional program that reinforces xenophagic capacity ^16–19^. Despite this powerful host defense system, successful intracellular pathogens have evolved strategies to circumvent host xenophagy and its regulation by TFEB. These evasion mechanisms typically involve interfering with pathogen recognition ^20,21^, blocking xenophagosomes formation ^22,23^ and its maturation into xenolysosomes ^24,25^, or impairing TFEB function by preventing its nuclear translocation ^14,26,27^, downregulating its expression ^28^, inhibiting its activation ^29^, or facilitating its degradation ^30^. A clearer understanding of these microbial tactics is essential for decoding host-pathogen dynamics and developing effective host-targeted therapies.

To unravel the complexity of host-pathogen interactions and microbial immune evasion strategies, well-characterized infection model systems are routinely used as experimental frameworks. Among these, *Lm* stands out as an effective and versatile model for dissecting xenophagy, owing to its distinct and well-characterized intracellular lifestyle. *Lm* is a Gram-positive, facultative intracellular bacterium that rapidly transitions from a vacuolar compartment to the host cytosol, a principal site of xenophagic surveillance. This transition is mediated by its major virulence determinants, LLO (encoded by *hlyA* gene) ^31–33^, a cholesterol-dependent pore-forming toxin, in coordination with accessory factors phospholipases C (PLCs), PI-PLC (encoded by *plcA* gene) and PC-PLC (encoded by *plcB* gene) ^34–36^, which together facilitate efficient vacuolar membrane disruption. Once in the cytosol, *Lm* not only survives but thrives by exploiting host actin dynamics. Its surface protein actin assembly-inducing protein A (ActA), encoded by *actA* gene, nucleates actin polymerization, initially forming dense actin clouds and subsequently elongating into polarized actin comet tails ^37–41^. These structures propel the bacterium through the cytosol and drive its spread into adjacent cells via protrusion and internalization, enabling tissue-level dissemination ^42–44^. Beyond intracellular motility, studies indicate that ActA also functions as a molecular shield, helping the bacterium evade detection by the xenophagic machinery, by preventing ubiquitination ^20,21,45^. This precisely coordinated transition from initial vacuolar entry to its subsequent evasion within the cytosol makes *Lm* an invaluable system for investigating host responses to both vacuole-contained and cytosol-exposed bacteria.

Despite significant insights into the role of LLO in promoting vacuolar escape, its broader impact on host xenophagy remains incompletely understood. While LLO-mediated membrane damage is known to trigger host autophagy ^46,47^, the precise mechanism by which *Lm* subsequently evades xenophagic degradation potentially through LLO remains unclear ^48^. Although the recruitment of the autophagy marker LC3 to intracellular bacteria has been documented, the critical distinction between xenophagy initiation and effective bacterial degradation is often underappreciated. This distinction is essential since LC3 recruitment alone may indicate an activated autophagic response, but without corresponding bacterial clearance, it may reflect an incomplete or abortive host defense.

To address this important gap, our study investigates the role of LLO in xenophagy evasion. We show that intracellular LLO binds LC3 in the host cytosol via a specific LIR motif, thereby preventing it from participating in xenophagy. In parallel, LLO disrupts lysosomal functionality and suppresses TFEB nuclear translocation, functionally crippling the autophagy-lysosomal pathway. Importantly, our findings demonstrate that pharmacological reactivation of TFEB or targeted disruption of the LLO-LC3 interaction effectively restores xenophagic clearance, resulting in enhanced elimination of the pathogen within host cells. Altogether, our study delineates a novel mechanism by which *Lm* subverts xenophagy, establishing LLO as a dual-function virulence factor that orchestrates both vacuolar escape and suppression of host degradative pathways.

## Results

### *Lm* evades xenophagy across intestinal epithelial and macrophage cells

To investigate the interplay between host xenophagy and survival strategies adopted by *Lm*, an *in vitro* infection model was established using cell lines of different tissue origin. These include human intestinal epithelial cells (HCT116), human monocytic cells (THP-1) and murine macrophage cells (RAW 264.7) (Fig. S1a). The intracellular replication kinetics of *Lm*, along with its ability to exploit host actin dynamics were analyzed. *Lm* demonstrated a strikingly accelerated exponential replication across different host cell types - HCT116 (Fig. S1b), THP1 (Fig. S1c) and RAW 264.7 (Fig. S1d) between 1- and 6-hours post infection (hpi), with a pronounced surge observed during 3 to 6h window. Simultaneously, analysis of host actin recruitment (Fig. S1, a and e) revealed a progressive rise in the proportion of *Lm* displaying actin clouds and actin tails over time. By 6hpi, more than 40% of intracellular *Lm* were associated with host actin structures across all tested cell lines. Live cell imaging at 6hpi captured the dynamic actin-based motility of *Lm* in HeLa cells (Supplementary movie 1). Bacterial movement occurred at a mean speed of ∼ 0.2µm/s. Individual bacteria were initially surrounded by rounded actin “clouds,” which then rapidly polarized into characteristic actin “comet tails,” propelling intracellular movement and enabling direct cell-to-cell spread. These observations underscore the remarkable adaptability of *Lm* across diverse host cell types and highlight the critical role of actin-mediated motility in facilitating its dissemination, especially during the later stages of infection. Building on these observations, we next investigated whether *Lm* undergoes xenophagy-mediated degradation by evaluating the involvement of autophagy and lysosomal machinery at two distinct time points: an early stage (3hpi) and a late stage (6hpi). These time points were chosen based on the observed rapid intracellular proliferation occurring between 3 and 6hpi. The association of autophagy markers LC3 and GABARAP, along with the lysosomal marker LAMP1, with the bacterial surface was assessed using fluorescence microscopy. This approach enabled visualization of xenophagosomes, indicated by LC3 or GABARAP recruitment to *Lm*, and xenolysosomes, marked by the co-recruitment of LC3 and LAMP1 to *Lm* (Fig. S1f). Infection in HCT116 (Fig. 1, a and b), THP-1 (Fig. 1, c and d), and RAW 264.7 (Fig. 1, e and f) cells revealed that *Lm* is targeted by xenophagy, as evidenced by its sequestration into LC3-positive vesicles at 3hpi. These xenophagic vesicles comprise both xenophagosomes and xenolysosomes, indicating progression toward lysosomal fusion. Quantification showed that ∼ 15-20% of *Lm* in HCT116 cells and 20-30% in macrophage cells were associated with xenophagic vesicles at this time point. By 6hpi, a clear decline in xenophagic targeting was observed across all cell types, reflected by a reduction in the percentage of xenophagosomes to ∼ 10-12%. This decrease was even more pronounced for xenolysosomes, which dropped to around 5-7%. These findings point to a progressive decline in both formation of xenophagosomes and their subsequent maturation into xenolysosomes over time. Similarly, the recruitment of GABARAP, an isoform of the mammalian autophagy-related protein 8 family, to *Lm* in THP-1 and RAW 264.7 cells showed marked decline over time. At 3hpi, ∼ 25-35% of bacteria were associated with GABARAP-positive structures. By 6hpi, this proportion dropped significantly to 10-15%, as shown in Fig. 1, g and h. Interestingly, we observed notable cell line-specific differences in autophagy and lysosomal marker recruitment. GABARAP recruitment to *Lm* remained undetected in HCT116 cells throughout the infection period (Fig. S2a), as opposed to the clear recruitment observed in THP-1 and RAW 264.7 cells. This suggests that a predominantly LC3-driven xenophagic response operates in HCT116 cells. While LAMP1 was clearly recruited to *Lm* in both HCT116 and THP-1 cells, no such recruitment was evident in RAW 264.7 cells at either time point (Fig. S2b), indicating a complete blockade in xenophagosomes maturation into degradative xenolysosomes in this macrophage model. Overall, the observed decline in the percentage of xenophagosomes and xenolysosomes across different host cell types suggests that *Lm* employs evasion strategies to counteract xenophagy-mediated clearance. By restricting xenophagosomes formation and potentially preventing their maturation into xenolysosomes, *Lm* effectively circumvents its degradation within the host.

**Figure 1.**
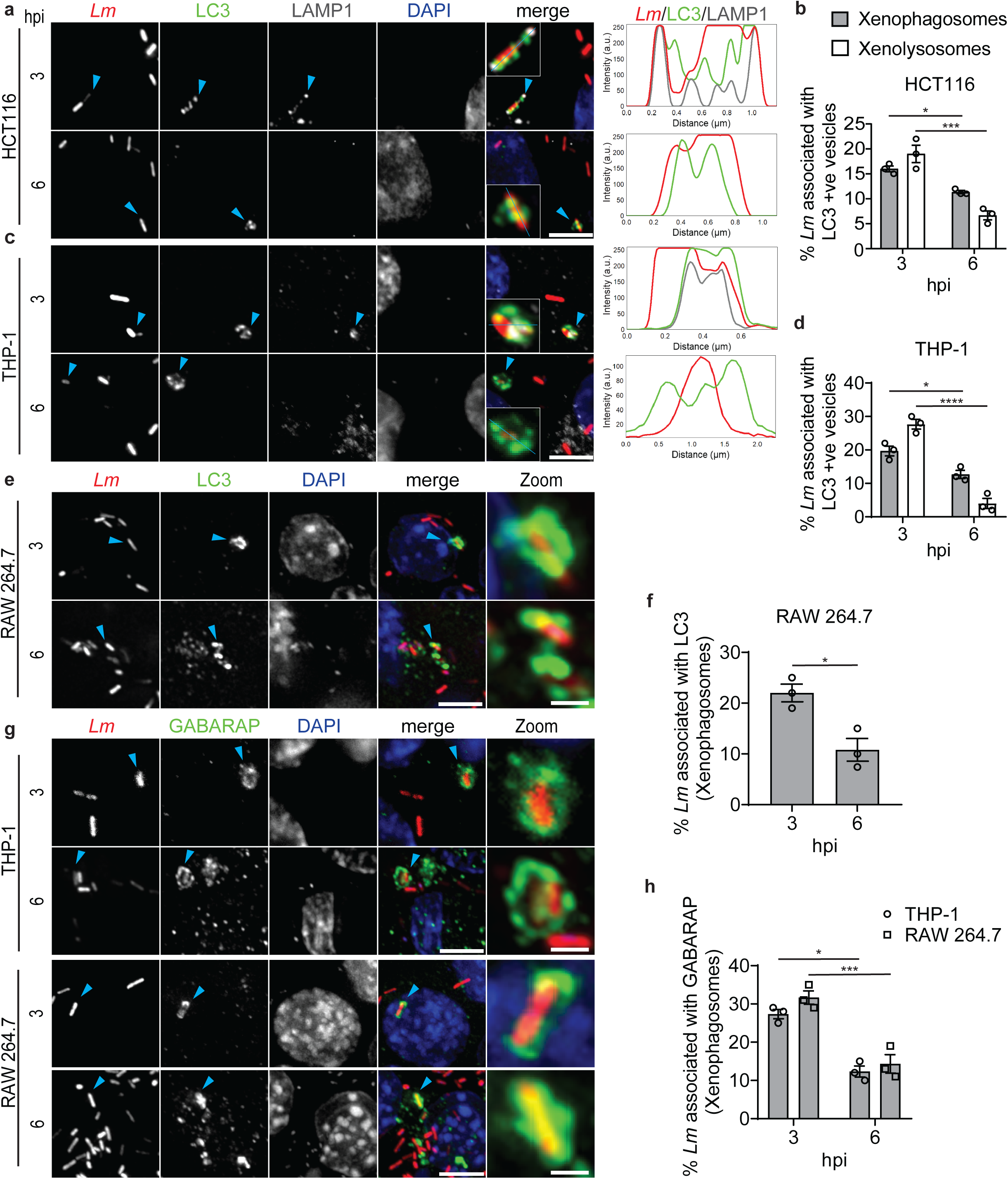
*Lm* evades xenophagy across intestinal epithelial and macrophage cells. The human intestinal cell line HCT116, differentiated human macrophage cell line THP-1 and mouse macrophage cell line RAW 264.7 were infected with mCherry expressing WT strain of *Lm* 10403S. At 3 and 6hpi, cells were immunostained using LC3, GABARAP, and LAMP1 antibodies to assess xenophagy. **a)** Fluorescence micrographs of HCT116 cells show xenophagosomes (recruitment of LC3 to *Lm*) and xenolysosomes (recruitment of both LC3 and LAMP1 to *Lm*) indicated by blue color arrowheads. Plot profiles along the blue line traced within the insets display intensity overlap of *Lm*, LC3, and LAMP1. **b)** Percentage of xenophagosomes and xenolysosomes at 3 and 6hpi in HCT116 cells. **c)** Fluorescence micrographs and plot profile of infected THP-1 cells show xenophagosomes and xenolysosomes. **d)** Percentage of xenophagosomes and xenolysosomes at 3 and 6hpi in THP-1 cells. **e)** Fluorescence micrographs show xenophagosomes in RAW 264.7 cells. **f)** Percentage of xenophagosomes at 3 and 6hpi in RAW 264.7 cells. **g)** Fluorescence micrographs of infected THP-1 and RAW 264.7 show recruitment of GABARAP to *Lm*. **h)** Percentage of xenophagosomes (recruitment of GABARAP to *Lm)* in THP-1 and RAW 264.7 cells at 3 and 6hpi. Scale bars: 5µm; zoom panel: 2µm. **b, d, f, h)** Data represents mean ± SEM (*N* = 3 independent experiments, *n* ≥ 100 bacteria per sample for each experiment). *P < 0.05, ***P < 0.001, ****P < 0.0001 using unpaired two-tailed t-test.

### Cytosolic (WT and Δ*actA*) and vacuolar *Lm* (Δ*hlyA*) show distinct fates and degradation patterns in intestinal epithelial cells

ActA and LLO, encoded by *actA* and *hlyA* genes respectively, are key *Lm* virulence factors crucial for its pathogenesis ^49^. Several studies suggest their involvement in modulating autophagy. For instance, ActA has been reported to potentially help *Lm* evade autophagic recognition by promoting actin polymerization on the bacterial surface ^20^. Conversely, LLO promotes phagosomal damage, allowing bacterial escape from the host cell vacuole, an event known to trigger autophagy activation ^47,50^. Despite these proposed roles, the precise mechanisms by which ActA and LLO influence autophagy are still not fully understood. To delve deeper into how autophagy differentiates between cytosolic and vacuolar *Lm*, we examined wild type (WT), Δ*actA* (lacking the *actA* gene), and Δ*hlyA* (lacking the *hlyA* gene) strains. Our experimental strategy was guided by prior studies showing that WT and Δ*actA* strains typically escape into the host cytosol within 30-60 minutes post-entry, whereas the Δ*hlyA* strain remains confined within the vacuole due to its inability to disrupt the vacuolar membrane ^51^. Leveraging these differences in intracellular localization, we assessed xenophagy by monitoring LC3 and LAMP1 recruitment to WT, Δ*actA*, and Δ*hlyA* strains in HCT116 at 3 and 6hpi using fluorescence microscopy. Gene deletions in the Δ*actA* and Δ*hlyA* strains were confirmed by colony PCR (Fig. S2d), and their extracellular growth kinetics, assessed via growth assays (Fig. S2e), showed no significant differences compared to the wild-type strain. Our results indicate that at 3hpi, both WT and Δ*actA* were targeted by xenophagy, with xenophagosomes (Fig. 2, a and b) and xenolysosomes (Fig. 2, a and c) each comprising ∼15-20% of the bacterial population. However, by 6hpi, xenophagy-associated compartments declined significantly in both strains, with Δ*actA* displaying a more pronounced reduction than WT. Since Δ*hlyA* lacks LC3 engagement, this negates the possibility of xenophagic or LC3-associated phagocytosis (LAP) degradation. However, the potential for Δ*hlyA* to be sequestered within LAMP1-positive compartments remains. To further characterize this mode of bacterial containment, we quantified LAMP1-only recruitment, defined as LAMP1-positive compartments lacking LC3 association. At 3hpi, ∼ 58% of Δ*hlyA* resided within LAMP1-positive compartments, whereas the percentages were notably lower for WT (∼20%) and Δ*actA* (∼5%). Over time, LAMP1-positive bacterial localization progressively declined in WT and Δ*actA* but remained stable in Δ*hlyA* (Fig. 2d). Additionally, Δ*hlyA* infected cells exhibited a substantially greater number of LAMP1 puncta relative to WT and Δ*actA* infected cells at 3 (Fig. S3a) and 6hpi (Fig. S3b). Due to observed differences in LAMP1 puncta number in WT, Δ*actA*, and Δ*hlyA* infected cells, we proceeded to analyze LC3 and LAMP1 colocalization events, which are indicative of autolysosomal number (Fig. 2e). Consistent with LAMP1 puncta quantification, Δ*hlyA* displayed significantly increased colocalization compared to WT and Δ*actA* at 3 (Fig. 2f) and 6hpi (Fig. 2g). The elevated number of LAMP1 puncta and LC3-LAMP1 colocalization events in Δ*hlyA* infected cells suggests a heightened lysosomal and autophagy response compared to WT, Δ*actA* infected and uninfected cells. Given the temporal decline in xenophagic vesicles in WT and Δ*actA* strains, we investigated the recruitment of the autophagy receptor p62 to *Lm* (Fig. S3c) to assess cargo recognition. At 3hpi, ∼ 15% of WT and ∼ 18% of Δ*actA* exhibited p62 recruitment; however, the percentage of p62-positive *Lm* decreased significantly at 6hpi to ∼ 4 to 7% for both strains (Fig. S3d). In contrast, Δ*hlyA* showed minimal p62 recruitment, consistent with its LC3 recruitment profile, suggesting diminished recognition by the xenophagy machinery due to its retention within vacuolar compartments (Fig. S3d). To explore the functional consequence of this differential host response to *Lm* strains, we compared intracellular bacterial load among WT, Δ*actA*, and Δ*hlyA* at 6hpi in infected HCT116, THP-1, and RAW 264.7 cells. In line with its extensive sequestration within LAMP1-positive compartments, Δ*hlyA* burden was significantly reduced in all cell types, indicating impaired intracellular survival, potentially due to phagolysosomal-mediated degradation. Compared to the WT strain, Δ*actA* showed similar intracellular load within THP-1 (Fig. S3f) and RAW 264.7 (Fig. S3g) cells but exhibited enhanced proliferation in HCT116 cells (Fig. S3e). This cell type-dependent disparity suggests that innate immune mechanisms such as inflammation and reactive oxygen species production ^52,53^ may contribute to limiting Δ*actA* proliferation in macrophages than in epithelial cells. Collectively, these findings highlight distinct autophagy-related outcomes for cytosolic and vacuolar populations of *Lm*. Although, WT and Δ*actA* strains escape into the cytosol and are initially targeted by xenophagy, this response wanes over time. Conversely, Δ*hlyA* strain appears to be primarily routed through the phagolysosomal pathway rather than being cleared via xenophagy or LAP. The pronounced autophagy and lysosomal response observed in Δ*hlyA* infected cells, coupled with its consistently reduced intracellular burden across multiple cell types, support a potential role for LLO in evading autophagy-lysosomal defenses and promoting *Lm* intracellular survival.

**Figure 2.**
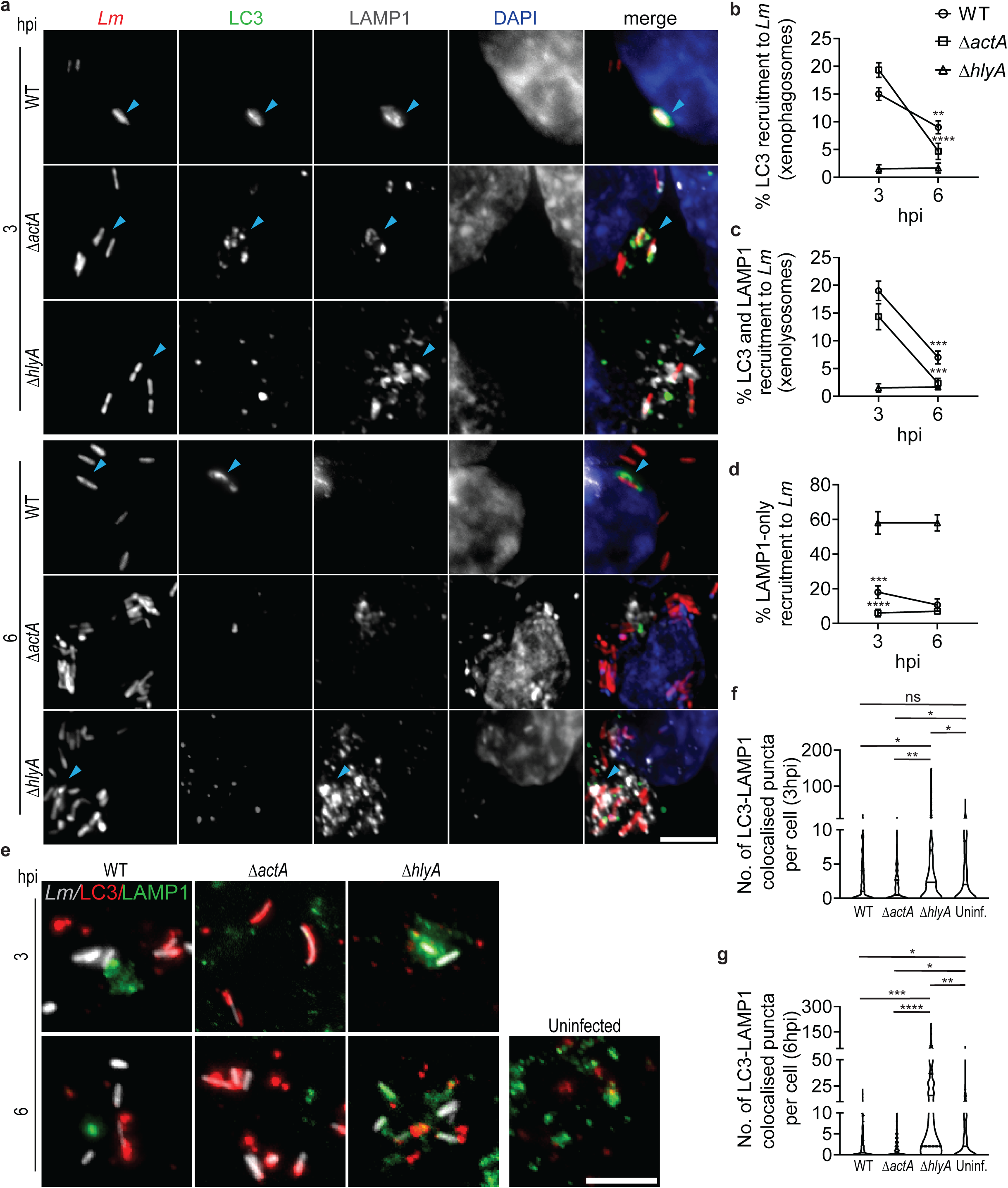
Cytosolic (WT and Δ*actA*) and vacuolar *Lm* (Δ*hlyA*) show distinct fates and degradation patterns in intestinal epithelial cells. HCT116 cells were infected with mCherry expressing WT, Δ*actA*, and Δ*hlyA* strains of *Lm* and immunostained with LC3 (green) and LAMP1 (grey) antibodies at 3 and 6hpi. **a)** Fluorescence micrographs of infected with WT, Δ*actA*, and Δ*hlyA* show xenophagosomes (recruitment of LC3 to *Lm*), xenolysosomes (recruitment of both LC3 and LAMP1 to *Lm*) and LAMP1-only recruitment to *Lm* (LAMP1 positive, LC3 negative *Lm*), indicated by blue color arrowheads. Percentage of xenophagosomes **b)**, xenolysosomes **c)**, and LAMP1-only recruitment to *Lm* in cells infected with WT, Δ*actA*, and Δ*hlyA* strains **d)**. **e)** Fluorescence micrographs and puncta colocalization between LC3 and LAMP1 in cells infected with WT, Δ*actA*, and Δ*hlyA* at 3 **f)** and 6hpi **g)**; uninfected cells serve as a control. Scale bars: 5µm. **b-d)** Data represents mean ± SEM (*N* = 3 independent experiments, *n* ≥ 100 bacteria per sample for each experiment). **P < 0.01, ***P < 0.001, ****P < 0.0001 using two-way ANOVA with Tukey’s multiple comparison test. **f, g)** Violin plots represent the density distribution of the number of autolysosomes at 3 and 6hpi in infected cells (*N* = 3 independent experiments, *n* ≥ 30 infected cells per sample for each experiment). The black bar indicates the interquartile range (IQR). Individual data points are shown as dots. *P < 0.05, **P < 0.01 using one-way ANOVA with Tukey’s multiple comparison test.

### *Lm* perturbs lysosomal acidification and hydrolytic activity to evade lysosomal-dependent degradation pathways in an LLO-dependent manner

Based on the observed differences in LAMP1 puncta abundance, LC3-LAMP1 colocalization, and LAMP1 recruitment to *Lm* in HCT116 cells infected with WT, Δ*actA*, and Δ*hlyA* strains, we sought to determine whether these variations reflect underlying differences in lysosomal function. Considering that lysosomal dysfunction impairs autophagy and disrupts other cellular processes dependent on lysosomal function, potentially leading to defects in intracellular recycling and homeostasis ^54^, we proceeded to evaluate lysosomal functionality. We assessed lysosomal acidity and proteolytic capacity in infected HCT116 cells. Lysosomal acidity was determined using LysoTracker staining ^55^ (Fig. 3a), a method that relies on selective accumulation of this dye in acidic organelles to serve as an indicator of lysosomal pH. To evaluate lysosomal proteolytic activity, we performed the DQ-BSA assay (Fig. 3c), wherein fluorescence is emitted upon enzymatic degradation of dye-quenched bovine serum albumin by lysosomal hydrolases ^56^. Quantification of LAMP1 and LysoTracker-positive *Lm* revealed that a significantly higher proportion (∼55%) of Δ*hlyA* bacteria localized within acidic, LAMP1-positive compartments compared to WT and Δ*actA* at 6hpi (Fig. 3b). Consistently, a greater fraction of Δ*hlyA* bacteria colocalized with both DQ-BSA and LysoTracker signals, indicating that ∼ 40% of Δ*hlyA* resided in hydrolytically active lysosomal compartments by 6hpi, whereas WT and Δ*actA* were largely negative for these markers (Fig. 3d). To assess overall lysosomal functionality, we quantified LysoTracker (Fig. 3e) and DQ-BSA (Fig. 3f) fluorescence intensities in infected HCT116 cells at 3 and 6hpi. Both indicators were significantly elevated in Δ*hlyA* infected cells compared to those infected with WT and Δ*actA*, reflecting increased lysosomal acidification and proteolytic activity. Interestingly, Δ*actA* infected cells displayed a marked reduction in both signals, especially at 6hpi, when compared to WT infected cells. Collectively, these results indicate that LLO expression dampens lysosomal acidification and enzymatic activity, thereby facilitating evasion of lysosomal-mediated degradation by WT and Δ*actA* strains.

**Figure 3.**
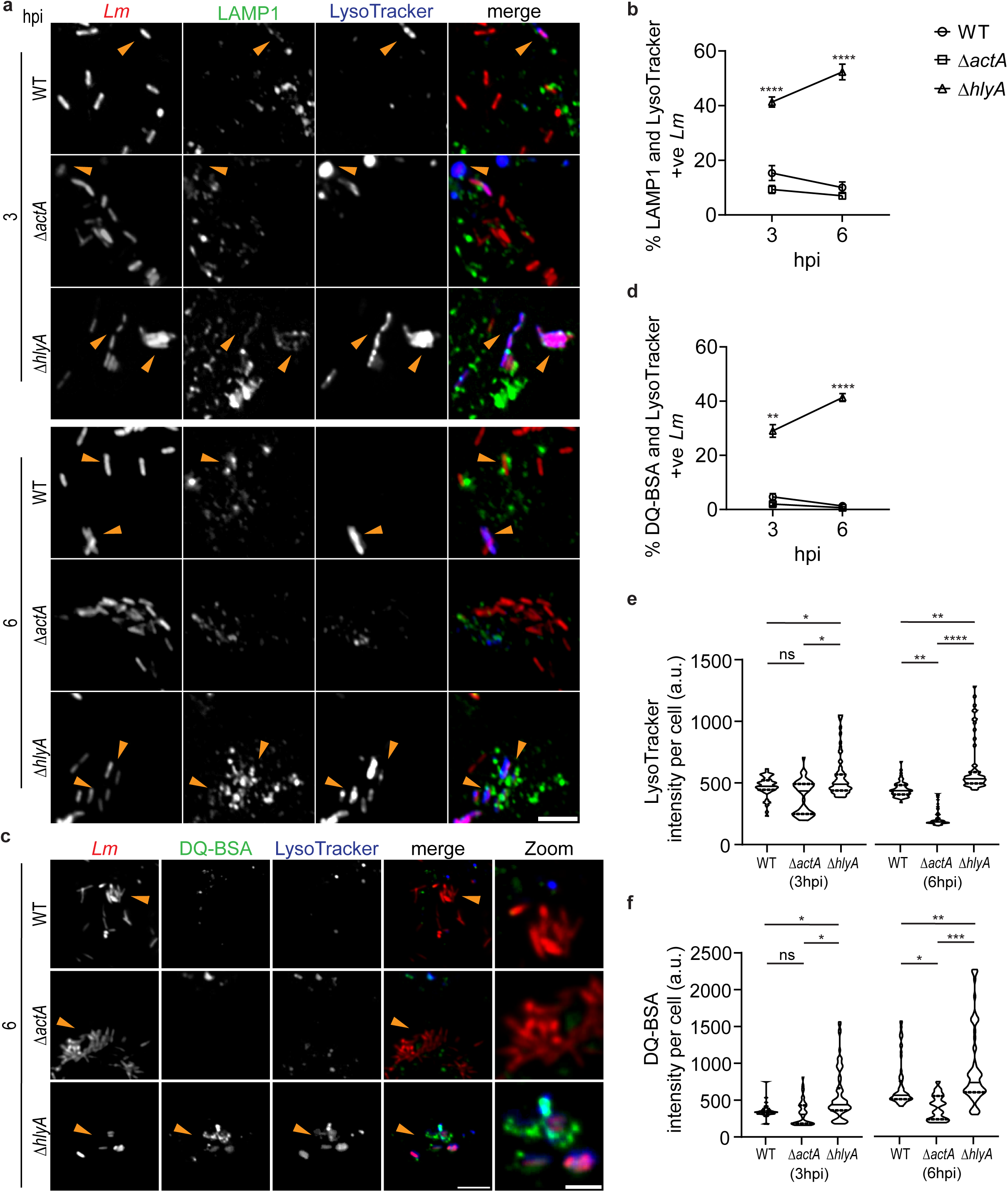
*Lm* perturbs lysosomal acidification and hydrolytic activity to evade lysosomal-dependent degradation pathways in an LLO-dependent manner. HCT116 cells were infected with mCherry expressing WT, Δ*actA*, and Δ*hlyA* strains of *Lm* and immunostained with LAMP1 antibody at 3 and 6hpi. Lysosomal acidification and degradative capacity were quantified using LysoTracker and DQ-BSA dyes. **a)** Fluorescence micrographs, and **b)** quantification of percentage LAMP1 and LysoTracker positive *Lm* in cells infected with WT, Δ*actA*, and Δ*hlyA* at 3 and 6hpi. **c)** Fluorescence micrographs, and **d)** quantification of percentage DQ-BSA and LysoTracker positive *Lm* in HCT116 cells infected with WT, Δ*actA* and Δ*hlyA* at 3 and 6hpi. Quantification of fluorescence intensity of LysoTracker **e)** and DQ-BSA **f)** at 3 and 6hpi in cells infected with WT, Δ*actA*, and Δ*hlyA*. Scale bars: **a)** 5µm, **c)** 10µm. **b, d)** Data represents mean ± S.E.M (*N* = 3 independent experiments, *n* ≥ 100 bacteria per sample for each experiment). ****P < 0.0001 using two-way ANOVA with Tukey’s multiple comparison test. **e, f)** Violin plots represent the density distribution of fluorescence intensity of LysoTracker and DQ-BSA at 3 and 6hpi in infected HCT116 cells (*N* = 3 independent experiments, *n* ≥ 30 infected cells per sample for each experiment). The black bar indicates the interquartile range (IQR). Individual data points are shown as dots. **P < 0.01, ***P < 0.001, ****P < 0.0001, ns > 0.9999 using one-way ANOVA with Tukey’s multiple comparison test.

### *Lm* suppresses TFEB nuclear translocation through the expression of LLO

Since *Lm* strains expressing LLO display impairment in both autophagy and lysosomal pathways, we hypothesized that these effects may be mediated through modulation of TFEB, a master transcriptional regulator of genes governing autophagy and lysosomal pathways. Under basal conditions, TFEB is phosphorylated and retained in the cytoplasm. Upon exposure to stress signals such as nutrient deprivation or intracellular infection, it becomes dephosphorylated and translocates to the nucleus to activate transcriptional programs linked to autophagy and lysosomal biogenesis ^57^. To determine whether TFEB is modulated in response to *Lm* infection, we assessed its activation by monitoring its dephosphorylation status and nuclear localization. Dephosphorylation was evaluated by immunoblotting, utilizing the phosphorylation-dependent mobility shift, whereby phosphorylated TFEB migrates more slowly and appears as a higher molecular weight “supershifted” band on the immunoblot ^58^. As a positive control, cells treated with Earle’s Balanced Salt Solution (EBSS), a known inducer of TFEB activation via starvation ^59^, exhibited complete conversion of phosphorylated TFEB to its dephosphorylated form, clearly visualized as a mobility shift on the immunoblot. WT and Δ*actA* infected cells displayed significantly reduced levels of dephosphorylated TFEB. In contrast, Δ*hlyA* infected cells showed significantly elevated levels of dephosphorylated TFEB relative to WT, Δ*actA*, and uninfected controls (Fig. 4, a and b), indicating enhanced TFEB activation in the absence of LLO. Consistent with TFEB results, LAMP1 level was significantly diminished in WT and Δ*actA* infected cells at 6hpi, whereas Δ*hlyA* infected cells exhibited increased LAMP1 levels (Fig. 4, a and c). The mCherry band in the immunoblot (Fig. 4a) reflects the presence of mCherry-labeled *Lm* in infected cells, with its intensity serving as a proxy for intracellular bacterial load. These findings reinforce a mechanistic link between TFEB suppression and compromised lysosomal function during infection with LLO expressing *Lm* strains. Additionally, we assessed TFEB nuclear translocation by quantifying nuclear TFEB intensity in infected HCT116 cells at 6hpi (Fig. 4d). Notably, Δ*hlyA* infected cells exhibited significantly higher TFEB nuclear localization compared to WT and Δ*actA* strains (Fig. 4e), indicating that in the absence of LLO, TFEB activation is not suppressed, thereby enabling robust autophagic and lysosomal response. To validate the direct effect of LLO on autophagy regulation, HCT116 cells were treated with 20µM purified recombinant LLO (rLLO), a concentration chosen based on assays demonstrating hemolytic activity (Fig. S4b) while maintaining host cell viability (Fig. S4a). The impact of rLLO on TFEB dephosphorylation (Fig. 5f) and LC3 I to LC3 II conversion (Fig. 5h) was examined by immunoblotting. In the presence of Torin1, an mTOR inhibitor that typically induces TFEB activation, rLLO treatment significantly reduced levels of dephosphorylated TFEB (Fig. 5g). Furthermore, rLLO exposure led to an increased LC3 II to I ratio following chloroquine (CQ) treatment (Fig. 5i) consistent with autophagosome accumulation due to impaired lysosomal fusion. Notably, LLO protein levels remained constant across Torin1- and CQ-treated samples (Fig. 5j and S4c), suggesting that LLO may be relatively resistant to autophagic degradation rather than being efficiently targeted as a cargo. Intriguingly, fluorescence microscopy revealed that intracellular LLO in infected cells appeared as dense punctate signals, persisting even at later time point (Fig. S4d). This was particularly evident in Δ*actA* infected cells, where bacteria escape from the primary vacuole but remain restricted to the initial host cell due to impaired actin-based motility, thereby preventing cell-to-cell spread and secondary vacuole formation ^60^. Despite this confinement, the sustained presence of dense LLO signal in Δ*actA* mutants implies its cytosolic relevance, even in the absence of its classic phagosomal function. Notably, Δ*actA* infected cells exhibited higher LLO levels per cell compared to WT (Fig. S4e), which coincided with a greater bacterial burden and a more pronounced suppression of autophagy, lysosomal function, and TFEB nuclear translocation. However, quantification of intracellular LLO levels in bacterial lysates revealed comparable expression between WT and Δ*actA* strains (Fig. S4, f and g), suggesting that the elevated cellular LLO levels observed in Δ*actA* infected cells likely result from more intracellular bacterial load rather than inherent differences in LLO production. Collectively, these findings highlight the pivotal role of LLO in intracellular survival of *Lm* by suppressing TFEB nuclear translocation and consequently blocking xenophagy.

**Figure 4.**
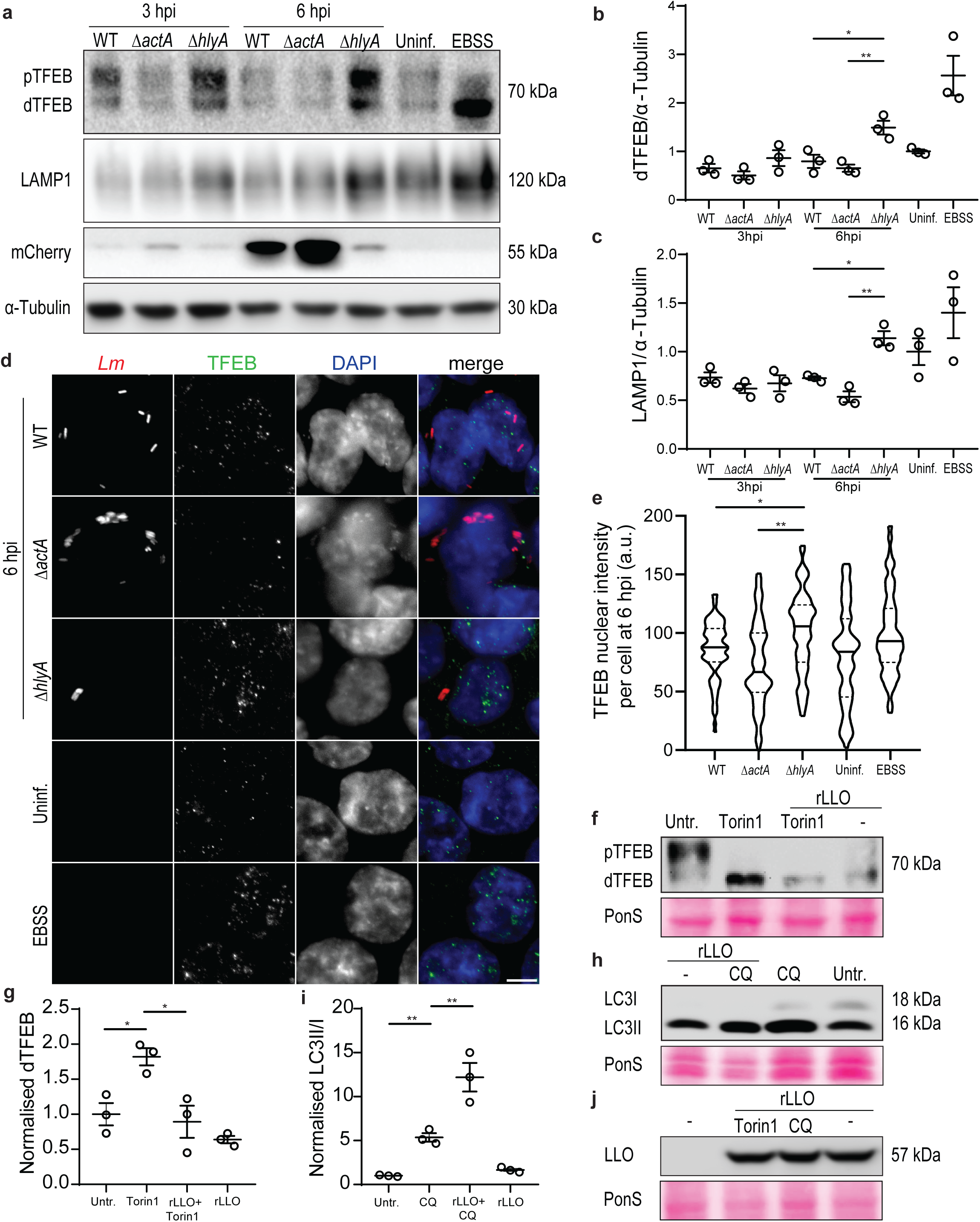
*Lm* suppresses TFEB nuclear translocation through the expression of LLO. **a)** Western blots depicting phospho- and dephosphorylated (faster-migrating TFEB band corresponds to the dephosphorylated form of the protein levels of TFEB), along with LAMP1 and mCherry levels in HCT116 cells following infection with WT, Δ*actA*, or Δ*hlyA* strains at 3 and 6hpi. Uninfected and EBSS-treated samples serve as controls. **b)** Dephosphorylated TFEB, and **c)** LAMP1 levels were quantified. **d)** Fluorescence micrograph, and **e)** quantification of nuclear fluorescence intensity of TFEB in HCT116 cells upon infection with WT, Δ*actA*, and Δ*hlyA* strains at 6hpi. HCT116 cells were treated with 20µM rLLO for 2h, with or without Torin1. Untreated cells and those treated only with Torin1 serve as controls. **f)** Western blots show phosphorylated and dephosphorylated TFEB. **g)** Dephosphorylated TFEB levels were quantified. HCT116 cells were treated with 20µM rLLO for 2h, with or without CQ. Untreated cells and CQ-only treated cells serve as controls. **h)** Western blots show lipidated (LC3II) and non-lipidated (LC3I) forms of LC3. **i)** LC3II/I ratio was quantified. **j)** Western blots display LLO levels in untreated cells and in cells treated with LLO alone, LLO combined with Torin 1, or LLO combined with CQ. Scale bar: 5µm. **b, c)** Data represents mean ± S.E.M (*N* = 3 independent experiments). *P < 0.05, **P < 0.01 using one-way ANOVA with Tukey’s multiple comparison test. **e)** Violin plot represents the density distribution of nuclear fluorescence intensity of TFEB per infected cell at 6hpi (*N* = 3 independent experiments, *n* ≥ 50 infected cells per sample for each experiment). The black bar indicates the interquartile range (IQR). Individual data points are shown as dots. *P < 0.05, **P < 0.01 using one-way ANOVA with Tukey’s multiple comparison test. **g, i)** Data represents mean ± S.E.M (N = 3 independent experiments). *P < 0.05, **P < 0.01 using one-way ANOVA with Tukey’s multiple comparison test.

**Figure 5.**
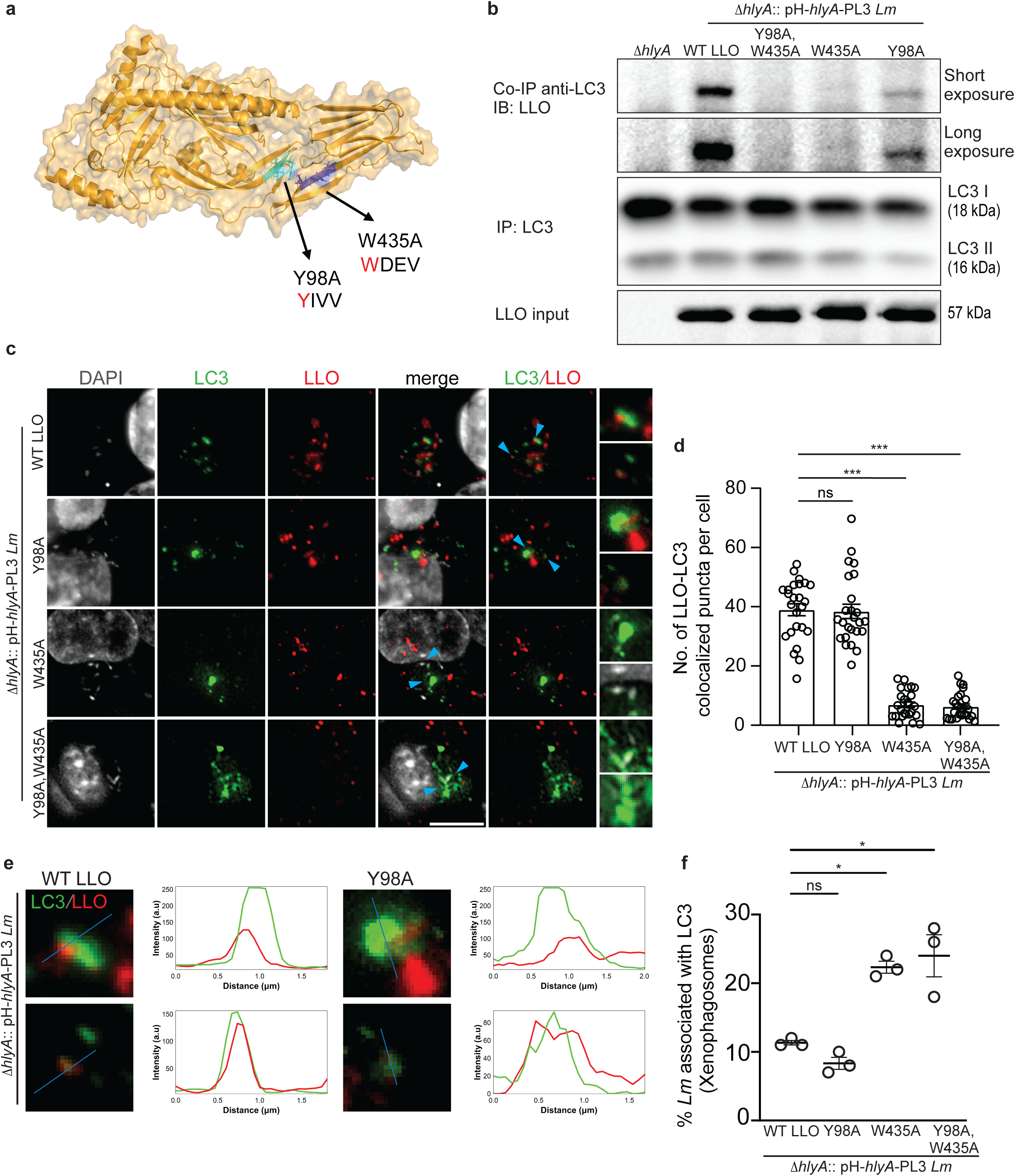
LLO binds to LC3 via an LIR motif and suppresses autophagy. **a)** The structure of monomeric LLO with two labeled putative LIR motifs, ^98^YIVV^101^ (located in the D2 domain) and ^435^WDEV^438^ (located in the D4 domain). Highlighted hydrophobic amino acids Y (tyrosine) and W (tryptophan) were mutated to A (alanine) using a site-directed mutagenesis approach to assess the role of the putative LIR motifs in LC3 interaction. **b)** LLO was obtained from the supernatant of bacterial cultures, (Δ*hlyA*, Δ*hlyA*::pH-*hlyA-*PL3, Δ*hlyA*::pH-*hlyA*(Y98A)-PL3, Δ*hlyA*::pH-*hlyA*(W435A)-PL3 and Δ*hlyA*::pH-*hlyA*(Y98A,W435A)-PL3) and is tested for its interaction with LC3 in HCT116 cell lysate. This interaction is examined using co-immunoprecipitation (Co-IP), via an antibody specific to LC3. **c)** Fluorescence micrograph and **d)** quantification of colocalization between LLO and LC3 in HCT116 cells infected with different strains as mentioned in **b**. **e)** Fluorescence micrographs of HCT116 cells infected with Δ*hlyA*::pH-*hlyA-*PL3 and Δ*hlyA*::pH-*hlyA*(Y98A)-PL3 strains show LLO and LC3 colocalization at 6hpi. Plot profiles reveal intensity overlap between LLO and LC3 along a defined blue line. **f)** Quantification of percentage xenophagosomes in HCT116 infected with Δ*hlyA*::pH-*hlyA-*PL3, Δ*hlyA*::pH-*hlyA*(Y98A)-PL3, Δ*hlyA*::pH-*hlyA*(W435A)-PL3 and Δ*hlyA*::pH-*hlyA*(Y98A,W435A)-PL3 strains. Scale bar: 5µm. **d)** Data represents mean ± S.E.M (*N* = 3 independent experiments, *n* ≥ 25 infected cells per sample for each experiment). ***P < 0.001, ns > 0.9999 using one-way ANOVA with Tukey’s multiple comparison test. **f)** Data represents mean ± S.E.M (N = 3 independent experiments, *n* ≥ 50 bacteria per sample for each experiment) *P < 0.05, **P < 0.01, ns > 0.9999 using one-way ANOVA with Tukey’s multiple comparison test.

### LLO binds to LC3 via an LIR motif and suppresses autophagy

Persisting in the host cytosol as a dense punctate signal, LLO coincides with suppression of TFEB nuclear translocation, along with impaired autophagy and lysosomal function. These findings led to the hypothesis that LLO, in addition to mediating vacuolar escape, may exert a direct role in subverting host degradative pathways. To explore the potential interaction between LLO and LC3, we analyzed the LLO protein sequence using the Eukaryotic Linear Motif (ELM) ^61^ and iLIR prediction ^62^ tools. This *in silico* analysis identified two putative LIR motifs: one located at residues 98-101 (^98^YIVV^101^) and another at residues 435-438 (^435^WDEV^438^) (Fig. 5a). To determine the functional importance of these motifs, site-directed mutagenesis was carried out to replace the conserved hydrophobic residue with alanine, resulting in the generation of three mutant *Lm* strains: Δ*hlyA*::pH-*hlyA*(Y98A)-PL3, Δ*hlyA*::pH-*hlyA*(W435A)-PL3 and Δ*hlyA*::pH-*hlyA*(Y98A,W435A)-PL3. No significant differences were observed in the extracellular growth kinetics (Fig. S5a) or in the levels of intracellular (Fig. S5, b and c) and secreted (Fig. S5, b and d) LLO between the mutant strains and the complemented control strain Δ*hlyA*::pH-*hlyA*-PL3 expressing WT LLO. To determine whether these mutations affect the interaction between LLO and LC3, co-immunoprecipitation (Co-IP) was performed. Culture supernatants from *Lm* strains expressing either WT or mutant LLO were incubated with lysates from HCT116 cells, followed by immunoprecipitation using an anti-LC3 antibody to capture LC3-bound LLO. Co-IP results revealed that LLO secreted from the Δ*hlyA*::pH-*hlyA*(Y98A)-PL3 mutant retained its ability to interact with LC3, comparable to the complemented Δ*hlyA*::pH-*hlyA*-PL3 strain. In contrast, LLO from both Δ*hlyA*::pH-*hlyA*(W435A)-PL3 and the double mutant Δ*hlyA*::pH-*hlyA*(Y98A,W435A)-PL3 failed to co-immunoprecipitate with LC3, indicating that the ^435^WDEV^438^ motif is critical for mediating this interaction (Fig. 5b). To corroborate the Co-IP results in a cellular context, we performed immunofluorescence microscopy on HCT116 cells infected with the respective mutants and complemented *Lm* strains to quantify colocalization between LLO and LC3 (Fig. 5c and Fig. 5e). Both WT LLO and LLOY98A exhibited clear colocalization with LC3, whereas the LLOW435A and the double mutant LLOY98A,W435A showed minimal overlap with LC3 signals (Fig. 5d). Interestingly, in strains harboring the ^435^WDEV^438^ motif mutation, LC3 signal was found to colocalize with the bacteria (Fig. 5e), a pattern not observed in the complemented strain or the Δ*hlyA*::pH-*hlyA*(Y98A)-PL3 strain. Quantification of LC3 recruitment confirmed that both Δ*hlyA*::pH-*hlyA*(W435A)-PL3 and the double mutant Δ*hlyA*::pH-*hlyA*(Y98A,W435A)-PL3 exhibited significantly enhanced LC3 recruitment compared to the Y98A mutant and the strain expressing wild-type LLO (Fig. 5f). Consistent with this increased LC3 recruitment, lysosomal marker analysis revealed that LLO mutants lacking the ^435^WDEV^438^ motif, exhibited significantly higher percentage of LAMP1 and LysoTracker positive bacteria (Fig. S5, f and g). Supporting these observations, intracellular CFU assays demonstrated a marked reduction in bacterial survival for both Δ*hlyA*::pH-*hlyA*(W435A)-PL3 and Δ*hlyA*::pH-*hlyA*(Y98A,W435A)-PL3 strains compared to the wild-type LLO and Y98A mutant strains (Fig. S5e). These findings identify the ^435^WDEV^438^ motif as the main contributor for LLO-LC3 interaction, contributing significantly to xenophagy evasion and enhancing intracellular survival of *Lm*. Overall, these findings reveal that LLO executes multiple strategies to disrupt the host autophagy-lysosomal pathway. It binds to LC3 in the cytosol through a defined LIR motif, ^435^WDEV^438^, limiting its availability from engaging in xenophagy.

### Restoring TFEB nuclear translocation enhances intracellular degradation of *Lm*

Since cytosolic *Lm* uses multiple strategies to evade autophagy-lysosomal pathway, we sought to counteract these evasion mechanisms through targeted chemical modulation. We made use of small molecules known to modulate autophagy. Torin1 (2µM) and rapamycin (4µM), both mTOR inhibitors that promote TFEB nuclear translocation ^63–65^, were used to restore transcriptional regulation of autophagy-related genes. To evaluate the contribution of autophagy at different stages, we also included an early-stage autophagy inhibitor SBI-0206965, an ULK1 inhibitor (2µM) ^66^ and a late-stage inhibitor, CQ (40µM) which blocks autophagosome–lysosome fusion ^67^ and assessed their impact on *Lm* intracellular proliferation (Fig. 6a). To rule out any direct antibacterial effects of these compounds, we first performed extracellular growth curve, which showed no significant differences in growth rates compared to untreated controls (Fig. 6b), indicating that the compounds do not affect *Lm* replication directly. Subsequent intracellular CFU assays in HCT116, THP-1, and RAW 264.7 cells (Fig. 6c) revealed distinct outcomes based on the type of autophagy modulation. Treatment with torin1 or rapamycin significantly reduced bacterial burden relative to untreated controls, while CQ treatment resulted in a marked increase in intracellular CFU. SBI-0206965 also showed an upward trend of *Lm* proliferation, although the change was not statistically significant across all cell types. These findings demonstrate that manipulation of the autophagy-lysosomal axis strongly influences *Lm* intracellular survival. To further validate the role of TFEB in this process, we silenced TFEB in HCT116 cells and measured bacterial burden at 6hpi (Fig. 6d). Both WT (Fig. 6e) and Δ*actA Lm* (Fig. 6f) strains showed significantly enhanced intracellular survival upon TFEB knockdown, reinforcing the critical role of TFEB in restricting *Lm* replication. Altogether, these results emphasize the importance of TFEB and the autophagy-lysosomal pathway as key regulators of host defense system against cytosolic *Lm*.

**Figure 6.**
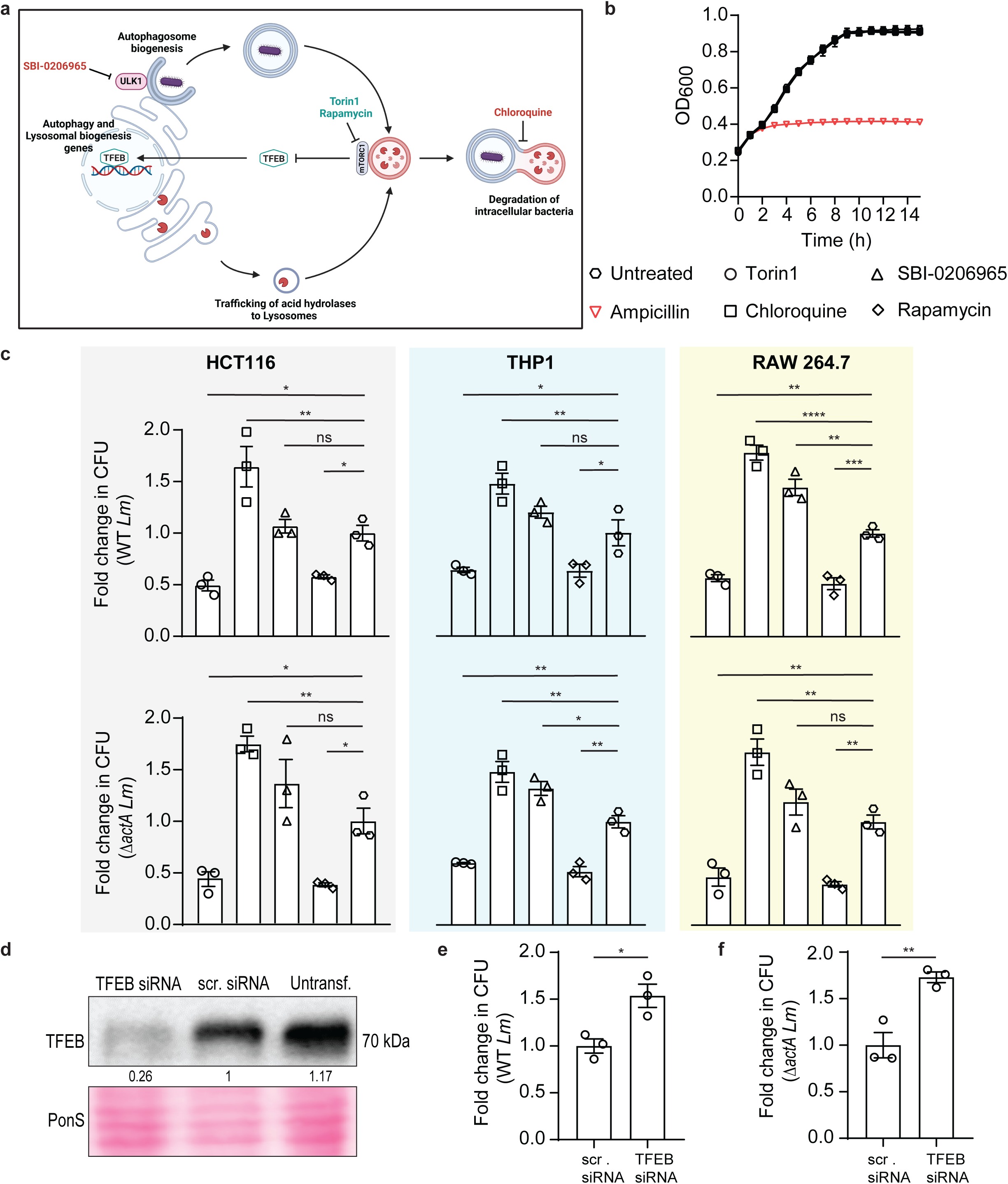
Restoring TFEB nuclear translocation enhances intracellular degradation of cytosolic *Lm*. **a)** Schematic illustrates the function of TFEB in autophagy and lysosomal biogenesis, highlighting its regulation via mTOR and the influence of small molecules including torin1, rapamycin, SBI-0206965, and CQ at various stages of these pathways. **b)** Growth curve of *Lm* following treatment with the small molecules mentioned in **a)**. Untreated samples and ampicillin-treated controls are included as controls. **c, d)** Quantification of intracellular bacterial load of WT and Δ*actA* in HCT116, THP-1, and RAW 264.7 cells at 6hpi. Infected cells were lysed, and colony-forming units (CFUs) were counted. CFU values from treated samples were normalized to those from untreated infected cells to determine fold change in bacterial burden. **e)** Western blot validation of TFEB siRNA knockdown efficiency. Intracellular load of **f**) WT and **g)** Δ*actA Lm* upon TFEB silencing. CFU data from siRNA-treated samples were normalized to CFU values from untreated infected cells to quantify fold change in bacterial burden. **c, d)** Data represents mean ± S.E.M (N = 3 independent experiments). *P < 0.05, **P < 0.01, ***P < 0.001, ****P < 0.0001, ns >0.9999 using one-way ANOVA with Dunnett’s multiple comparison test. **f, g)** Data represents mean ± S.E.M (N = 3 independent experiments). *P < 0.05, **P < 0.01 using unpaired two-tailed t-test.

## Discussion

*Lm* is known to deploy multiple strategies to evade xenophagic clearance, allowing it to persist within host cells. Among its virulence factors, ActA and Internalin K (InlK) have been shown to subvert xenophagy by hijacking host proteins. ActA promotes actin polymerization to shield the bacterium ^45^, while InlK recruits major vault protein (MVP) to mask its surface ^68,69^. Through these distinct yet convergent mechanisms, both ActA and InlK prevent xenophagic recognition and targeting. Additionally, one study reports that *Lm* modulates host lipid signaling through phospholipases C (PlcA and PlcB), reducing phosphatidylinositol 3-phosphate (PI3P) levels and stalling pre-autophagosomal structures, thereby interfering with autophagosome maturation ^70,71^. Building upon this established framework, our study reveals that *Lm* leverages a multipronged xenophagy disruptive function of its principal virulence factor, LLO. Beyond its canonical role in vacuolar escape, LLO simultaneously interferes with multiple components of the host autophagy-lysosomal axis. We show that intracellular LLO interacts with LC3 via a defined LIR motif, in the cytosol to reduce its availability to participate in the formation of xenophagosome. Simultaneously, LLO impairs lysosomal function and represses activation of the master regulator TFEB by inhibiting its nuclear translocation, further dampening the host xenophagy machinery. Crucially, our data reveals that pharmacological activation of TFEB or targeted disruption of the LLO-LC3 interaction is sufficient to restore xenophagy, leading to enhanced intracellular bacterial clearance. Taken together, our findings redefine the role of LLO as a critical effector that antagonizes xenophagic clearance post-vacuolar escape, expanding our understanding of immune evasion tactics of *Lm*.

Our comparative analysis of xenophagic targeting across distinct *Lm* strains differing in intracellular localization and motility: WT (cytosolic and spreading), Δ*actA* (cytosolic but non-spreading), and Δ*hlyA* (vacuolar) provides new insights into bacterial survival strategies. Both WT and Δ*actA* strains, successfully evade xenophagy at later stages, indicating that actin-based motility is not essential for avoiding xenophagic degradation. In contrast, the vacuole-confined Δ*hlyA* strain fails to engage with LC3 yet strongly associate with LAMP1, indicating its retention within phagolysosomes rather than diversion to either xenophagy or LAP pathway. These findings reinforce the centrality of LLO-mediated phagosomal escape and suggest that ActA is dispensable for intracellular survival. Our results, along with some of previous studies, demonstrate that ActA is not strictly required for intracellular survival ^50,68,72^, defined as the ability of *Lm* to remain viable and replicate within a host cell in the absence of ActA. In line with this, our finding and reports from others show comparable intracellular burden for both WT and Δ*actA* strains across multiple host cell types and tissues, including the colon and ileum ^43,73,74^, highlighting that actin-based motility is not universally required for intracellular survival. Nonetheless, ActA remains critical when considering bacterial dissemination beyond the initial site of infection. Its role in polymerization of actin enables *Lm* to traverse cellular barriers and colonize distal tissues like the brain and placenta ^43,75–77^. Thus, while dispensable in some contexts, ActA confers a selective advantage to *Lm* during systemic infection by enabling efficient dissemination to distant tissues, balancing the metabolic cost of actin polymerisation with the benefit of enhanced tissue tropism and pathogenesis.

In addition to evading xenophagic targeting, cytosolic *Lm* disrupts the abundance and functionality of degradative autophagosomal and lysosomal compartments, as evidenced by reduced autolysosomal population, altered lysosomal pH and activity. This impairment suggests that cytosolic *Lm*, independent of actin-based motility, also interferes with downstream autophagic maturation and lysosomal function. These observations are in line with previous reports implicating cholesterol-dependent cytolysins (CDCs) such as perfringolysin O (PFO), pneumolysin (PLY), and streptolysin O (SLO) to impair lysosomal integrity ^78–80^. The shared ability of these toxins underscores a conserved immune evasion strategy adapted by Gram-positive pathogens to enhance intracellular survival.

In parallel, our findings reveal that LLO suppresses activation of TFEB, a critical transcription factor responsible for autophagic and lysosomal biogenesis at 6hpi. While one study has demonstrated that TFEB translocates to the nucleus during the initial phase of *Lm* infection, this nuclear translocation diminishes by 4hpi, suggesting a tightly regulated temporal response. Although LLO-induced membrane damage has been identified as the upstream trigger for TFEB activation, the experimental assessments of TFEB nuclear translocation were limited to an early timepoint (1hpi) ^17^. This constraint precludes a full understanding of the temporal dynamics and regulatory mechanisms governing TFEB localization beyond the initial phase of infection. Indeed, multiple bacterial and viral pathogens have been reported to target TFEB as part of their host immune evasion tactics ^16^, further emphasizing its role as a key regulator of antimicrobial responses.

Central to our study is the mechanistic role of LLO in subverting host autophagy. While previous reports have suggested that LLO induces autophagy ^47^, this response likely represents a host-driven mechanism aimed at eliminating damaged vacuoles post-escape, rather than a bacterial survival tactic. A key observation from our data is the sustained secretion of LLO into the cytosol. Notably, we detect LLO as dense, punctate signals within the host cytosol at later stages of infection, prompting an investigation into its post-escape role. Mechanistically, we demonstrate that secreted LLO directly binds to LC3 via a functional LIR motif, ^435^WDEV^438^, with a preferential interaction likely toward the lipidated LC3 which is LC3II form. This interaction is supported by our findings of LC3 puncta formation and LC3II accumulation upon *Lm* infection or LLO treatment. However, the conjugation of LC3 to vesicles remains ambiguous, especially regarding the nature of these vesicles whether they exhibit single or double membranes, whether they are fully sealed during the LLO-LC3 interaction. Furthermore, it is unclear whether LLO exists freely within the cytosol or is associated with specific membrane-bound compartments. Defying the autophagic degradation of Cry5B toxin of *Bacillus thuringiensis* in *Caenorhabditis elegans* ^81^, LLO exhibits an extraordinary resilience, bypassing degradation even when autophagy is pharmacologically activated by Torin1. This persistent cytosolic accumulation suggests that the LLO-LC3 complex forms a stable, non-productive structure that evades degradative pathway. We propose that LLO acts as a molecular decoy, sequestering LC3II away from its role in xenophagosome formation. Reported evidence of LLO forming aggregates further supports this hypothesis ^82^. One report suggests these cytosolic LLO structures resemble neurodegeneration-associated aggregates ^83^ and another show LLO colocalizing with ubiquitin and p62 though whether these are subject to autophagic degradation remains unclear ^84^. Additionally, while degradation of LLO monomers and aggregates via the ubiquitin-proteasome system (N-end rule pathway) is well established ^31,85,86^, our data raise the possibility that some LLO aggregates may escape this route by entering autophagic vesicles and blocking fusion with lysosomes. Similar autophagy-evasion strategies have also been observed in other pathogens. For example, the non-structural small (NSs) protein from Rift Valley fever virus binds LC3 through a LIR motif, leading to nuclear retention of LC3 and xenophagy inhibition ^87^. Other examples include IcsB effector of *Shigella flexneri* binds ATG5 to prevent xenophagosome formation ^88^, RavZ protease of *Legionella pneumophila* that cleaves LC3 from autophagosomal membranes to prevent its autophagic degradation ^23^, and M2 protein of Influenza A virus hijacks LC3 to facilitate viral egress ^89^. Together, these examples underscore a shared evolutionary strategy among intracellular pathogens to mimic host short linear motifs (SLiMs) ^90,91^, particularly LIR motif to manipulate autophagy and ensure intracellular persistence. Understanding this molecular mimicry will reveal foundational principles in host-pathogen interactions and could accelerate the development of novel antimicrobial therapies ^92^.

Importantly, our study demonstrates that restoring TFEB activation using pharmacological activators or disrupting LLO-LC3 interaction via targeted mutagenesis, significantly reduces the intracellular burden of *Lm*. TFEB activation not only restores lysosomal function but also boosts LC3 and other autophagy proteins levels ^93,94^, producing sufficient LC3 and other factors to outcompete LLO for binding and thereby counteract its sequestration effect. Furthermore, the reduced intracellular burden of *Lm* following disruption of the LLO-LC3 interaction substantiates our conclusion that ActA is not necessary for xenophagy evasion.

Overall, these findings establish LLO not only as a pore-forming effector essential for vacuolar escape but also as a potent modulator of the host autophagy-lysosomal pathway. The dual targeting of LC3 and TFEB reveals parallel axes of immune subversion, reinforcing the idea that *Lm* has evolved a highly integrated approach to manipulating host cellular defenses. While the critical role of LLO in xenophagy inhibition is clearly established, questions remain regarding the spatial and temporal regulation of its activity and aggregation propensity within host compartments. Though its pore-forming function is pH-dependent ^95^, it is plausible that additional regulatory layers and differential expression ^96^ modulate its dual functionality, balancing immune evasion with host cell viability. Of particular interest is the mechanism by which LLO inhibits TFEB nuclear translocation, which remains undefined. Emerging studies have identified LC3 lipidation and GABARAP conjugation to selective autophagic membranes as key triggers of TFEB activation during lysophagy and xenophagy, respectively ^97,98^. The sequestration of LC3 by LLO may therefore interfere with this signaling axis thereby suppressing TFEB activation. Unravelling this regulatory crosstalk will be essential for fully characterizing multifunctionality of LLO and developing host-targeted therapeutic strategies to strengthen autophagic surveillance.

While the early phase (∼ 1hpi) of *Lm* infection is defined by the responses to listeriolysin O (LLO), including release of cathepsins from lysosomes ^79^ and TFEB translocation ^17^. However, the progressive decline of these responses despite the continued presence of LLO raises a critical question on the role of LLO at the later stages of infection. Our study addresses this question by investigating how sustained LLO exposure influences intracellular survival beyond the acute cellular response window. We demonstrate that *Lm* leverages LLO not only to breach vacuolar confinement but also to subvert host xenophagic defenses through LC3 binding, inhibition of TFEB nuclear translocation, and lysosomal disruption. This late-stage immune interference allows the pathogen to persist within host cells, revealing an intricately coordinated strategy for long-term survival. Given its central role, LLO stands out as a compelling therapeutic target, with multiple studies supporting its inhibition to effectively control *Lm* infection both *in vitro* and *in vivo* ^99–101^. These insights underscore the multifaceted impact of LLO on host-pathogen dynamics and reinforce the therapeutic potential of modulating autophagy-related regulators.

## Supplementary figures

**Figure S1.**
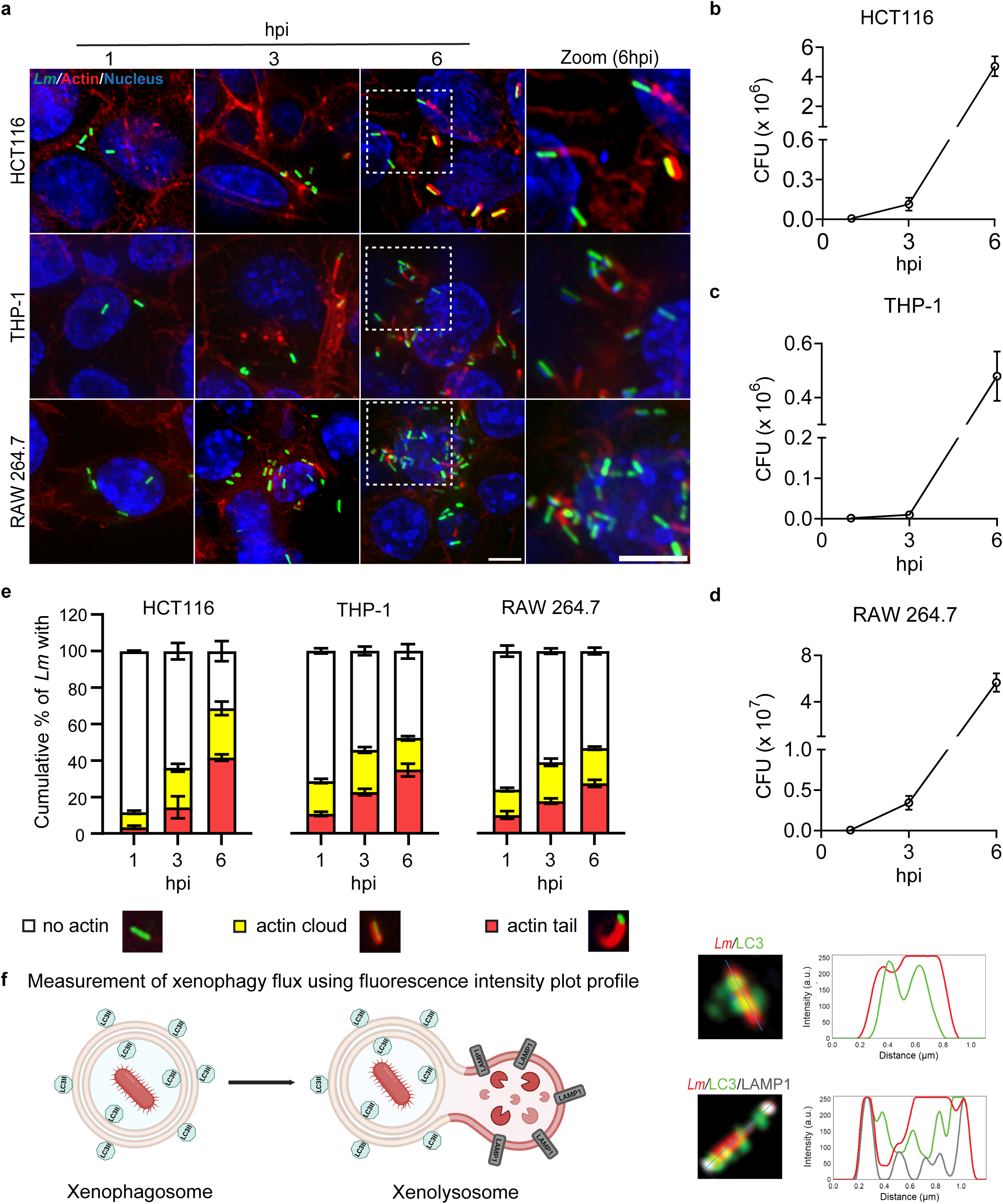
Intracellular *Lm* exhibits exponential growth and host actin-based motility at later time points of infection across different cells. HCT116, THP-1, and RAW 264.7 cells were infected with GFP expressing WT strain of *Lm*. Cells were stained with phalloidin dye to visualize host actin network at 1, 3, and 6hpi. **a)** Fluorescence micrographs depict intracellular proliferation of *Lm* and host actin-based clouds and tail formation by *Lm* across different time points. **b**-**d**) Intracellular growth kinetics of *Lm* in HCT116, THP-1, and RAW 264.7 cells. **e)** Quantification of *Lm*-mediated host actin cloud and actin tail formation at indicated timepoints in HCT116, RAW 264.7, and THP-1 cells. **f)** Schematic shows measurement of xenophagy using fluorescence intensity plot profile. Scale bars: 5µm; zoom panel: 5µm. **b, c, d)** Data represents mean ± S.E.M (*N* = 3 independent experiments). **e)** Data represents mean ± S.E.M (*N* = 3 independent experiments, *n* ≥ 100 bacteria per sample for each experiment).

**Figure S2.**
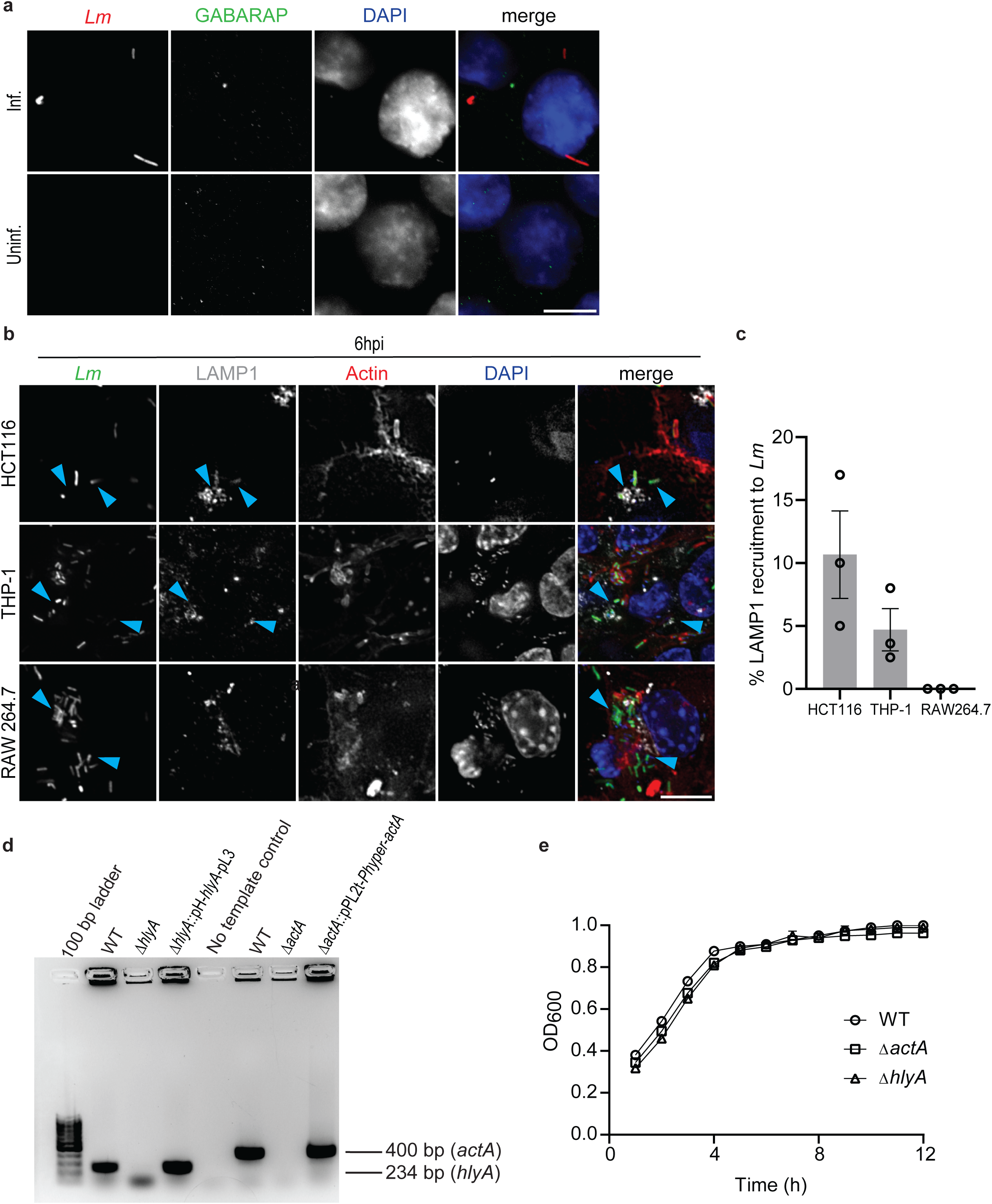
**a)** Representative fluorescence micrographs of infected HCT116 cells demonstrate the absence of GABARAP recruitment to *Lm*. **b)** Representative fluorescence micrographs of HCT116, THP-1, and RAW 264.7 cells infected with *Lm* show the recruitment of actin and LAMP1 to the bacteria. **c)** Quantitation of percentage LAMP1 recruitment to *Lm* at 6hpi. **d)** PCR confirmation of *hlyA* and *actA* gene deletions in putative Δ*hlyA* and Δ*actA* strains of *Lm*. Complementary strains Δ*hlyA*::pH-*hlyA*-PL3 and Δ*actA*::pPL2t.*P_hyper_*-*actA* served as controls. **e)** Growth curves of WT, Δ*actA*, and Δ*hlyA* strains in BHI broth at 37°C, assessing their proliferation over time. Scale bar: 5µm.

**Figure S3.**
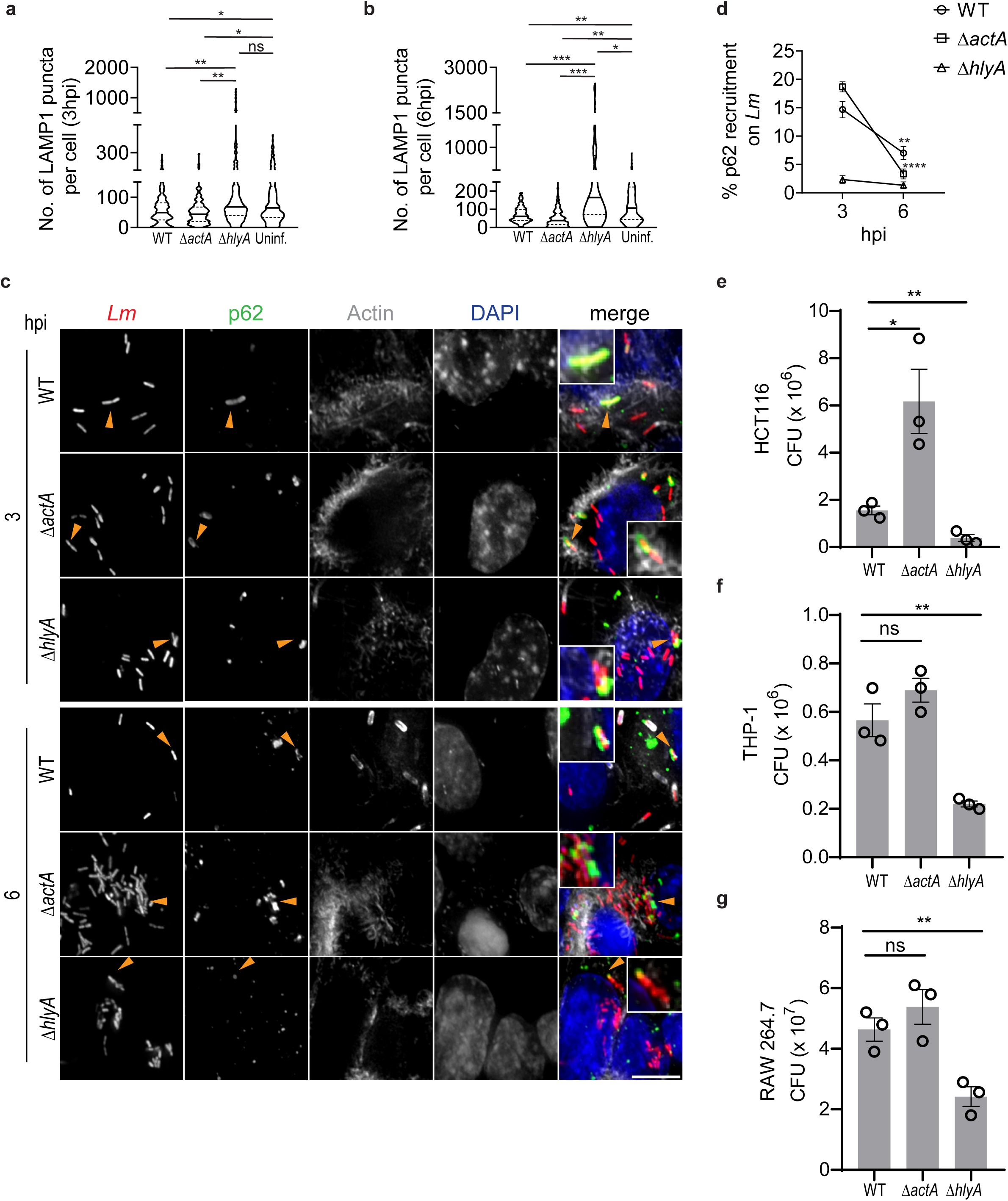
Quantification of the number of LAMP1 puncta in HCT116 cells infected with WT, Δ*actA*, and Δ*hlyA* at 3 **a)** and 6hpi **b)**, fluorescence micrographs are shown in Figure 2 **e**. HCT116 cells were infected with mCherry expressing WT, Δ*actA*, and Δ*hlyA* strains of *Lm* and immunostained with p62 (green) at 3 and 6hpi. Host actin network was labeled using phalloidin dye (grey). **c)** Fluorescence micrographs and **d)** quantification of percentage p62 recruitment to *Lm* at 3 and 6hpi in HCT116 cells. Quantification of intracellular CFU of WT, Δ*actA*, and Δ*hlyA* in HCT116 **e)**, THP-1 **f)**, and RAW 264.7 cells **g)** at 6hpi. Intracellular CFU was determined by lysing the infected cells, plating on BHI agar plate, and counting the CFUs. Scale bar: 5µm. **a, b)** Violin plots represent the density distribution of the number of lysosomes at 3 and 6hpi in infected HCT116 cells (*N* = 3 independent experiments, *n* ≥ 30 infected cells per sample for each experiment). The black bar indicates the interquartile range (IQR). Individual data points are shown as dots. *P < 0.05, ****P < 0.0001 using one-way ANOVA with Tukey’s multiple comparison test. **d)** Data represents mean ± S.E.M (*N* = 3 independent experiments, *n* ≥ 100 bacteria per sample for each experiment). **P < 0.01, ****P < 0.0001 using two-way ANOVA with Tukey’s multiple comparison test. **e-g)** Data represents mean ± S.E.M (N = 3 independent experiments). *P < 0.05, **P < 0.01, ns (not significant) >0.9999 using one-way ANOVA with Tukey’s multiple comparison test.

**Figure S4.**
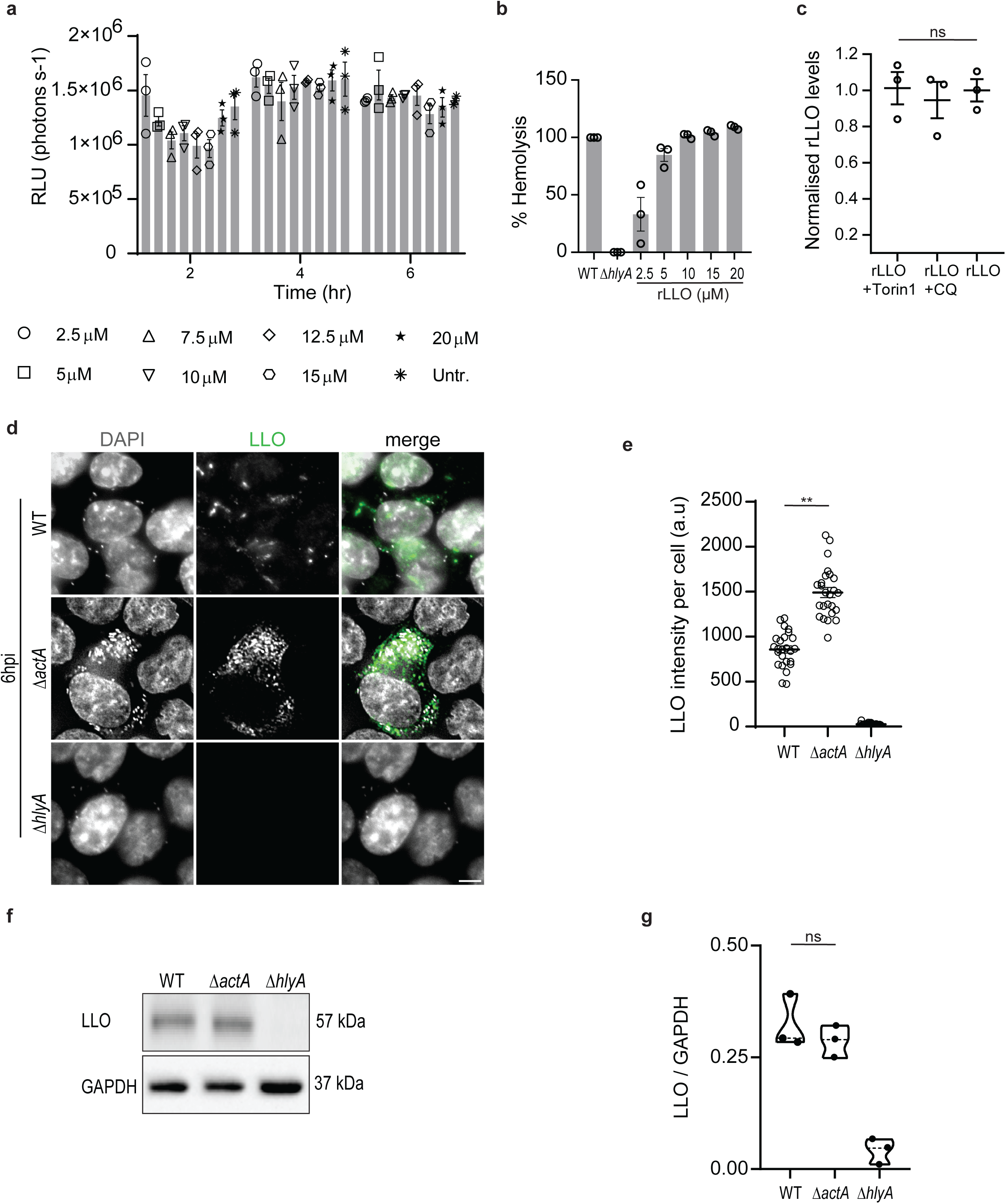
**a)** Cell viability assessment in HCT116 cells following rLLO treatment, quantified as relative luminescence units (RLU). **b)** Percentage hemolysis observed at varying concentrations of rLLO. **c)** Western blot-based quantification of LLO levels in HCT116 cells treated with 20µM rLLO for 2h, with and without Torin1 or CQ treatment (data corresponding to Figure 4j).**d)** Fluorescence micrographs and **e)** quantification of LLO fluorescence intensity in HCT116 cells infected with WT, Δ*actA*, and Δ*hlyA* strains at 6hpi. **f)** Western blot analysis and **j)** quantification of LLO protein levels in WT, Δ*actA*, and Δ*hlyA* strains of *Lm*. Scale bar: 5µm. **c, g)** Data represents mean ± S.E.M (*N* = 3 independent experiments), ns >0.9999, using one-way ANOVA with Tukey’s multiple comparison test. **e)** Data represents mean ± S.E.M (*N* = 3 independent experiments, *n* = 25 infected cells per sample for each experiment), **P < 0.01, using one-way ANOVA with Tukey’s multiple comparison test.

**Figure S5.**
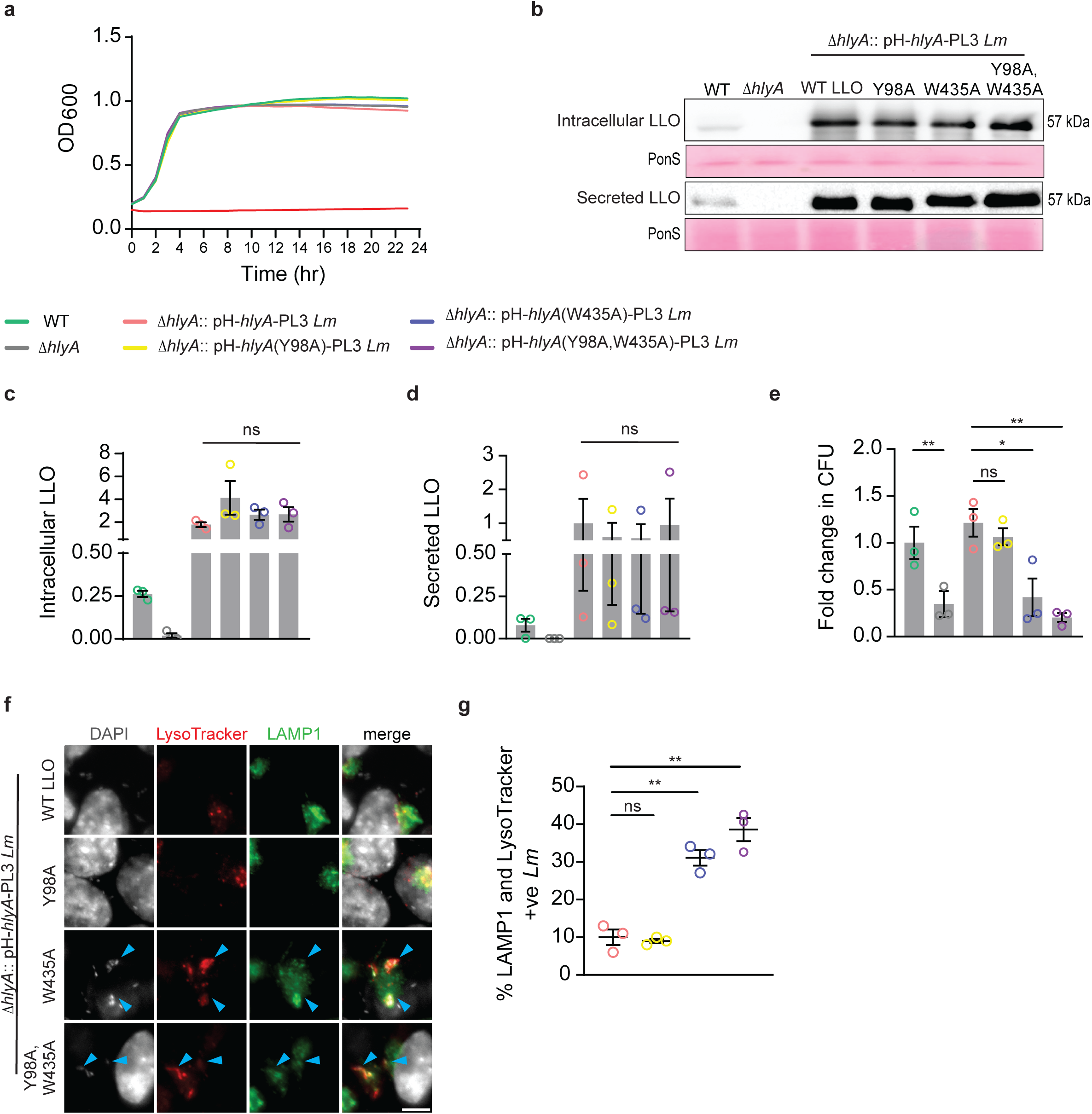
Disruption of LLO-LC3 interaction reinstates host xenophagy against *Lm*. **a)** Growth curve of WT, Δ*hlyA*, Δ*hlyA*::pH-*hlyA-*PL3, Δ*hlyA*::pH-*hlyA*(Y98A)-PL3, Δ*hlyA*::pH-*hlyA*(W435A)-PL3 and Δ*hlyA*::pH-*hlyA*(Y98A,W435A)-PL3 strains. **b-d)** Western blot analysis of intracellular and secreted LLO levels across the same strains. **e)** Intracellular CFU quantification of WT, Δ*hlyA*, Δ*hlyA*::pH-*hlyA-*PL3, Δ*hlyA*::pH-*hlyA*(Y98A)-PL3, Δ*hlyA*::pH-*hlyA*(W435A)-PL3 and Δ*hlyA*::pH-*hlyA*(Y98A,W435A)-PL3 strains in HCT116 cells at 6hpi. CFU values from WT *Lm* infected strains were normalized to those from cells infected with mutant and complemented strains to calculate the fold change in bacterial burden. **f, g)** Representative fluorescence micrographs, and quantification of percentage LAMP1 and LysoTracker positive *Lm* in HCT116 cells infected with Δ*hlyA*::pH-*hlyA*-PL3, Δ*hlyA*::pH-*hlyA*(Y98A)-PL3, Δ*hlyA*::pH-*hlyA*(W435A)-PL3 and Δ*hlyA*::pH-*hlyA*(Y98A,W435A)-PL3 strains at 6hpi. **b-d)** Data shown as mean ± S.E.M (N = 3 independent experiments), ns > 0.9999 using one-way ANOVA with Tukey’s multiple comparison test. **e)** Data represents mean ± S.E.M (N = 3 independent experiments). *P < 0.05, **P < 0.01, ns >0.9999 using one-way ANOVA with Dunnett’s multiple comparison test. **g)** Data represents mean ± S.E.M (N = 3 independent experiments, *n* ≥ 50 bacteria per sample for each experiment) *P < 0.05, **P < 0.01, ns > 0.9999 using one-way ANOVA with Tukey’s multiple comparison test.

**Supplementary movie 1: *Lm* live cell.avi**

Intracellular and intercellular host actin-mediated movement of *Lm*.

## Materials and Methods

### Bacterial strains and plasmids

All *Lm* strains used in this study (listed in Table 1) are derived from 10403S, a streptomycin-resistant isolate of strain 10403S (kind gift from Prof. Daniel A Portnoy, UCB, USA). *Lm* was cultured in Brain Heart Infusion (BHI) (BD Difco, 237500) broth to perform infection assays as follows: a single colony was inoculated into 2 mL of filter-sterilized (0.22µm) BHI broth and incubated overnight at 37°C in a tube lying flat. The resulting primary culture was diluted 1:10 into fresh BHI broth and grown at 37°C at 200 rpm until the optical density at 600 nm (OD_600_) reached 0.8–1.0, typically within 3-5h. This secondary culture was then used for infection assays in host cells. *E. coli* strains (listed in Table 2) were grown in Luria-Bertani (LB) (HiMedia, M1245) broth at 37°C. The following antibiotic concentrations were used: streptomycin (G-Biosciences, RC-197) (200μg/mL), chloramphenicol (Sigma-Aldrich, C0378) (7.5μg/mL for *Lm* and 10μg/mL for *E. coli*) and tetracycline (Sigma-Aldrich, 64-75-5) (2μg/mL). All plasmids used in this study are listed in Table 3. To perform bacterial imaging, plasmids pPL2-mCherry and pPL2-GFP were transconjugated from *E. coli* SM10 (kind gift from Prof. Daniel A Portnoy, UCB, USA) into *Lm* to generate mCherry and GFP-expressing *Lm* strains. pPL2t.*P_hyper_*-*actA* and pH-*hlyA-*PL3 plasmids were transconjugated from *E. coli* SM10 into Δ*actA* and Δ*hlyA Lm* strains respectively to generate complemented strains Δ*actA*::pPL2t.*P_hyper_*-*actA* and Δ*hlyA*::pH-*hlyA-*PL3 *Lm*. Colony PCR was used to verify these complemented strains with primers detailed in Table 3. Point mutations were introduced into *hlyA* gene within pH-*hlyA-*PL3 plasmid using SPRIP (Single-Primer Reactions IN Parallel) mutagenesis method ^102^. The primers used to create the specific nucleotide change are listed in Table 3. The mutations were confirmed through DNA sequencing. Further, pH-*hlyA-*PL3 plasmids carrying point mutation were transformed into competent *E. coli* SM10 and selected on a chloramphenicol containing LB plate. Donor *E. coli* SM10 containing the mutant plasmid and recipient Δ*hlyA Lm* were streaked in an overlapping manner on a non-selective BHI plate and incubated for 24h at 37°C. Transconjugants were then selected by picking overlapping colonies and plating them on a BHI plate containing streptomycin and chloramphenicol.

**Table 1.** Details of bacterial strains used.

| Bacterial strain | Description | Reference |
| --- | --- | --- |
| ***Listeria monocytogenes (Lm) 10403S*** |  |  |
| WT | Kind gift from Prof. Daniel A Portnoy, UCB, USA | <sup>104</sup> |
| $\Delta actA$ | Kind gift from Prof. Daniel A Portnoy, UCB, USA | <sup>105</sup> |
| $\Delta hlyA$ | Kind gift from Prof. Daniel A Portnoy, UCB, USA | <sup>106</sup> |
| WT mCherry | mCherry expressing *Lm* | This study |
| WT GFP | GFP expressing *Lm* | This study |
| $\Delta actA$ mCherry | mCherry expressing $\Delta actA$ *Lm* | This study |
| $\Delta hlyA$ mCherry | mCherry expressing $\Delta hlyA$ *Lm* | This study |
| $\Delta actA::pPL2t.P_{hyper-actA}$ | Complemented strain expressing WT *actA* gene to restore *actA* expression in $\Delta actA$ background | This study |
| $\Delta hlyA::pH-hlyA$-PL3 | Complemented strain expressing WT *hlyA* gene to restore *hlyA* expression in $\Delta hlyA$ background | This study |
| $\Delta hlyA::pH-hlyA$(Y98A)-PL3 | Complemented strain expressing *hlyA* gene carrying Tyr to Ala mutation at 98 amino acid position in $\Delta hlyA$ background | This study |
| $\Delta hlyA::pH-hlyA$(W435A)-PL3 | Complemented strain expressing *hlyA* gene carrying Trp to Ala mutation at 435 amino acid position in $\Delta hlyA$ background | This study |
| $\Delta hlyA::pH-hlyA(Y98A,W435A)$-PL3 | Complemented strain expressing *hlyA* gene carrying Tyr to Ala mutation at 98 amino acid position and Trp to Ala mutation at 435 amino acid position in $\Delta hlyA$ background | This study |
| ***E. coli* strain** |  |  |
| XL10-Gold Ultracompetent Cells | For transformation of pH-*hlyA*-PL3 carrying point mutations in *hlyA* gene following site-directed mutagenesis | This study |
| SM10 | Transconjugation donor strain (Kind gift from Prof. Daniel A Portnoy, UCB, USA) | <sup>107</sup> |
| SM10-mCherry | Transconjugation donor strain | This study |
| SM10-GFP | Transconjugation donor strain | This study |
| SM10- pPL2t.*P<sub>hyper</sub>-actA* | Transconjugation donor strain | This study |
| SM10- pH-*hlyA*-PL3 | Transconjugation donor strain | This study |
| SM10- pH-*hlyA*(Y98A)-PL3 | Transconjugation donor strain | This study |
| SM10- pH-*hlyA*(W435A)-PL3 | Transconjugation donor strain | This study |
| SM10- pH-*hlyA*(Y98A,W435A)-PL3 | Transconjugation donor strain | This study |

**Table 2.** Details of plasmids used.

| Plasmid | Description | Reference |
| --- | --- | --- |
| pPL2-mCherry | Integrative plasmid pPL2 carrying *mCherry* gene under a constitutive promoter with chloramphenicol resistance (Kind gift from Prof. Daniel A Portnoy, UCB, USA) | <sup>108</sup> |
| pPL2-GFP | Integrative plasmid pPL2 carrying *gfp* gene under a constitutive promoter with chloramphenicol resistance (Kind gift from Prof. Daniel A Portnoy, UCB, USA) | <sup>108</sup> |
| pPL2t.*P<sub>hyper</sub>-actA* | Shuttle integration vector for overexpression of *actA* gene from a constitutive HyPer promoter at *tRNA<sup>Arg</sup>* locus with tetracycline resistance (Kind gift from Prof. Daniel A Portnoy, UCB, USA) | <sup>109</sup> |
| pH-*hlyA*-PL3 | Shuttle integration vector for overexpression of *hlyA* gene from a constitutive HyPer promoter at *tRNA<sup>Arg</sup>* locus with chloramphenicol resistance (Kind gift from Prof. Daniel A Portnoy, UCB, USA) | <sup>109</sup> |
| pH-*hlyA*(Y98A)-PL3 | *hlyA* gene mutation (HlyA Y98A) results in the substitution of tyrosine with alanine at amino acid position 98 | This study |
| pH-*hlyA*(W435A)-PL3 | *hlyA* gene mutation (HlyA W435A) results in the substitution of tryptophan with alanine at amino acid position 435 | This study |
| pH-*hlyA*(Y98A,W435A)-PL3 | *hlyA* gene mutation (HlyA Y98A,W435A) results in the substitution of tyrosine with alanine at amino acid position 98 and substitution of tryptophan with alanine at amino acid position 435 | This study |
| Lifeact-mCherry | plasmid #193300 | Addgene |

**Table 3.**
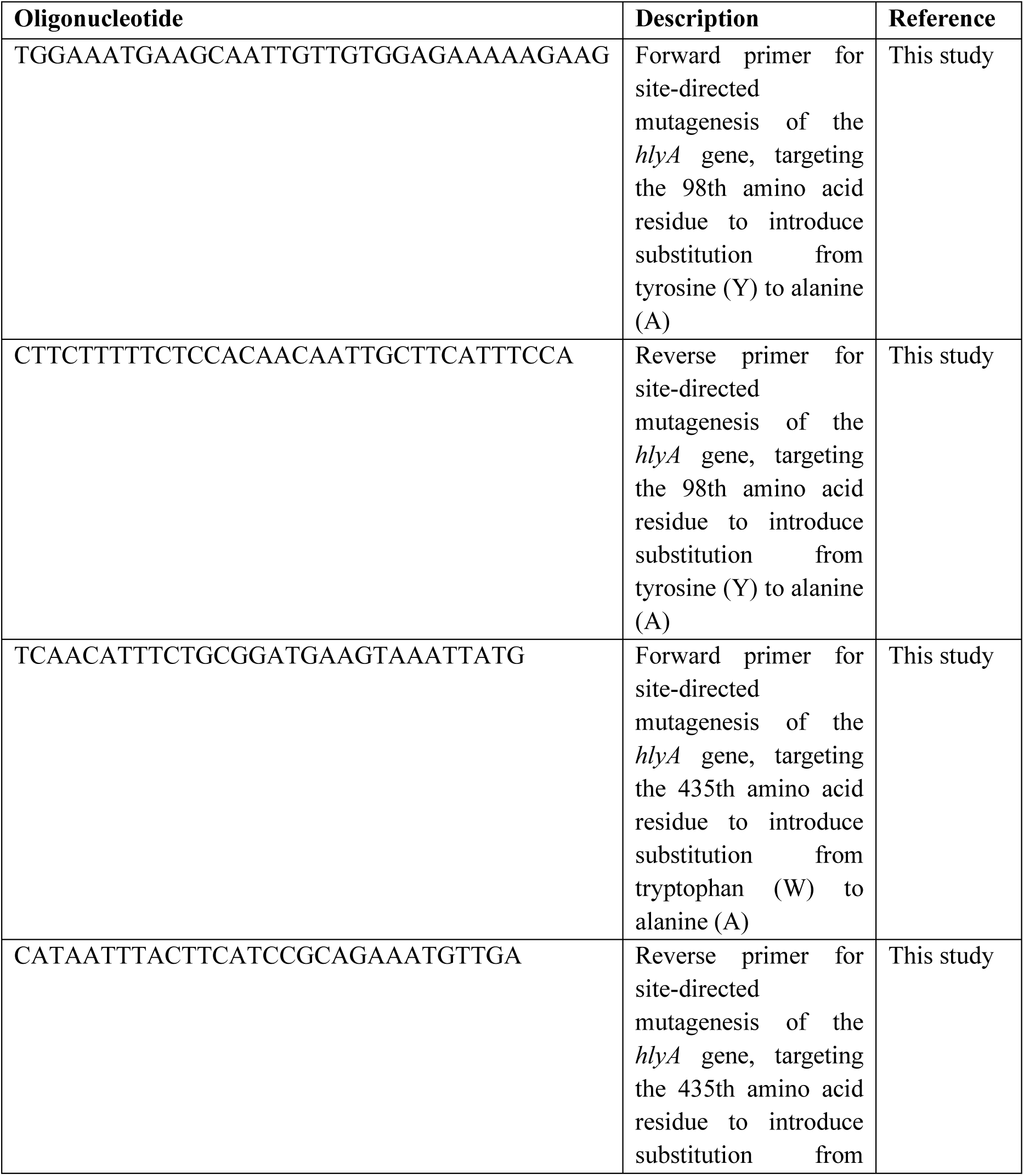
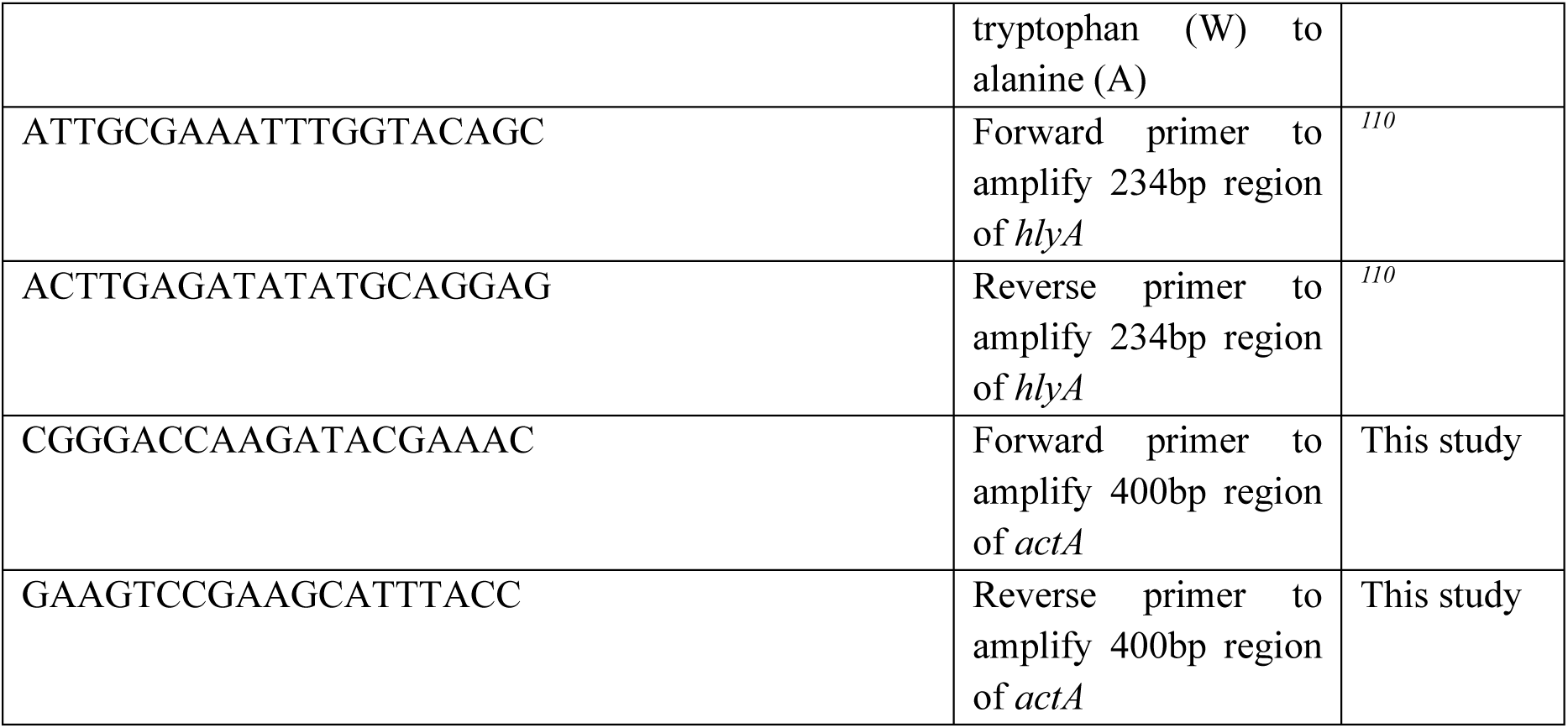
Details of oligonucleotides used.

### Cell culture

HCT116 cells were maintained in McCoy’s 5A medium (Sigma-Aldrich, D5648) supplemented with 2.2g/L sodium bicarbonate (Sigma-Aldrich, S5761), 100U/mL penicillin-streptomycin (Pen-Strep) (Gibco, 15140122), and 10% fetal bovine serum (FBS) (Gibco, 10270106). THP-1 cells were cultured in RPMI medium (Sigma-Aldrich, R6504) supplemented with 2.2g/L sodium bicarbonate, 100 U/mL Pen-Strep, and 10% FBS. RAW 264.7 and HeLa cells were grown in DMEM (Sigma-Aldrich, D5648) supplemented with 3.7g/L sodium bicarbonate, 100 U/mL Pen-Strep, and 10% FBS. All cell lines were incubated at 37°C in a 5% CO₂ humidified atmosphere and were procured from NCCS, Pune, India (Cell Repository). For infection assays, THP-1 cells were differentiated into macrophages by pre-treatment with 25ng/mL phorbol 12-myristate 13-acetate (PMA) (Sigma-Aldrich, P1585) for 24h. Following differentiation, cells were washed once with 1X DPBS (Sigma-Aldrich, D5773) and cultured in fresh RPMI medium for an additional 24h prior to use.

### Intracellular Colony Forming Unit (CFU) infection Assay

HCT116 cells were infected at a multiplicity of infection (MOI) of 50 for 1h. Differentiated THP-1 and RAW 264.7 cells were infected at an MOI of 20 and 5, respectively, for 30min. HeLa cells were infected at an MOI of 100 for 1h. Infections were performed in incomplete media (without FBS). Following bacterial incubation, cells were washed three times with 1X DPBS. To eliminate extracellular bacteria, gentamicin (50μg/mL) (Genticyn) was added to incomplete media for 20min. Cells were then washed with 1X DPBS and incubated in complete media containing 25μg/mL gentamicin to prevent extracellular bacterial growth, with or without the compound (listed in Table 4), until the time of collection. To lyse host cells and release intracellular bacteria, cells were treated with 0.1% ice-cold Triton X-100 (Millipore, 648466) in 1X DPBS. Lysates were plated on BHI agar and incubated at 37°C for 24h. Intracellular bacterial replication or survival was quantified by counting CFU.

**Table 4.** Details of antibodies, reagents and other resources used.

| *Resources* | *Procurement detail* | *Product ID* |
| --- | --- | --- |
| **Antibodies** |  |  |
| Mouse monoclonal anti-LC3 (IF and CO-IP) | MBL Life Science | M152-3 |
| Rabbit polyclonal anti-LC3 (IB) | Sigma-Aldrich | L7543 |
| Rabbit monoclonal anti-GABARAP | Abcam | ab109364 |
| Rabbit monoclonal anti-LAMP1 | Cell Signaling Technology | 9091 |
| SQSTM1/p62 | Abcam | ab56416 |
| Rabbit polyclonal anti-TFEB | Cell Signaling Technology | 4240 |
| Mouse monoclonal anti-RFP | ChromoTek | 6G6 |
| Rabbit polyclonal anti-Listeriolysin (LLO) | Abcam | ab200538 |
| Rabbit polyclonal anti-α-Tubulin | Cell Signaling Technology | 2144 |
| Rabbit monoclonal anti-GAPDH | Cell Signaling Technology | 14C10 |
| Anti-Mouse IgG-Atto 488 | Sigma-Aldrich | 62197 |
| Anti-Rabbit IgG-Atto 550 | Sigma-Aldrich | 43328 |
| Anti-Rabbit IgG-Atto 633 | Sigma-Aldrich | 41176 |
| Anti-rabbit IgG, HRP-linked | Cell Signaling Technology | 7074 |
| Anti-mouse IgG, HRP-linked | Cell Signaling Technology | 7076 |
| IF: Immunofluorescence, Co-IP: Co-immunoprecipitation, IB: Immunoblotting |  |  |

| **Reagents** |  |  |
| --- | --- | --- |
| Dulbecco's Modified Eagle Medium (DMEM) | Sigma-Aldrich | D5648 |
| McCoy's 5A Medium | Sigma-Aldrich | M4892 |
| RPMI-1640 Medium | Sigma-Aldrich | R6504 |
| Sodium bicarbonate | Sigma-Aldrich | S5761 |
| Sodium Pyruvate (100mM) | Gibco | 11360070 |
| Trypsin-EDTA Solution 10X | Sigma-Aldrich | 59418C |
| Fetal bovine serum (FBS) | Gibco | 10270106 |
| Live Cell Imaging Solution | Invitrogen | A59688DJ |
| Penicillin-Streptomycin (Pen-strep) (10,000U/mL) | Gibco | 15140122 |
| Dulbecco's Phosphate Buffered Saline (DPBS) | Sigma-Aldrich | D5773 |
| Earle's Balanced Salt Solution (EBSS) 10X | Sigma-Aldrich | E7510 |
| Lipofectamine 2000 | Invitrogen | 11668019 |
| Phorbol 12-myristate 13-acetate (PMA) | Sigma-Aldrich | P1585 |
| Brain Heart Infusion (BHI) broth | BD Difco | 237500 |
| Luria Bertani (LB) Broth | HiMedia | M1245 |
| Agar powder | HiMedia | GRM026 |
| Syringe Filter, Pore Size 0.22µm | Sartorius | 16532 |
| Triton X-100 | Millipore | 648466 |
| Protease inhibitor cocktail | Roche, Sigma-Aldrich | COEDTAF-RO |
| Phenylmethanesulfonyl fluoride (PMSF) | Sigma-Aldrich | P7626 |
| Ponceau S | Sigma-Aldrich | 6226-79-5 |
| CellTiter-Glo (CTG) | Promega | G7570 |
| Trichloroacetic acid (TCA) | Thermo Fisher Scientific | 76-03-9 |
| TFEB esiRNA | Sigma-Aldrich | EHU059261 |
| siRNA Universal Negative Control | Sigma-Aldrich | SIC001 |
| Dithiothreitol (DTT) | Thermo Fisher Scientific | R0862 |
| Cell Imaging Coverglass with 4 chambers | Eppendorf | 0030 742.028 |
| 96-well plate (for growth curve and hemolysis assay) | Corning | CLS3370 |
| 96-well plate (CTG assay) | Corning | 3917 |
| 24-well plate | CytoOne | CC7682-7524 |
| 12-well plate | CytoOne | CC7682-7512 |
| FluoroTrans PVDF (Polyvinylidene fluoride) transfer membrane | PALL Life Sciences | 79548A |
| Enhanced chemiluminescence substrate (ECL) | Clarity Bio-Rad | 170-5061 |
| Bradford Reagent | Sigma-Aldrich | B6916 |
| Cell scraper | Tarsons | 960051 |
| Recombinant LLO | Abcam | ab83345 |
| **Compounds** |  |  |
| Torin 1 | Tocris Bioscience | 4247 |
| Rapamycin | Sigma-Aldrich | 553210 |
| Chloroquine diphosphate salt | Sigma-Aldrich | C6628 |
| SBI-0206965 | Selleck Chemicals | S7885 |
| **Antibiotics** |  |  |
| Streptomycin Sulphate | G-Biosciences | RC-197 |
| Chloramphenicol | Sigma-Aldrich | C0378 |
| Tetracycline hydrochloride | Sigma-Aldrich | 64-75-5 |
| Ampicillin sodium salt | Sigma-Aldrich | A9518 |
| Gentamicin | Genticyn | NA |
| **Dyes** |  |  |
| Phalloidin–Atto 633 | Sigma-Aldrich | 68825 |
| BODIPY FL Conjugate of BSA (Green) | Sigma-Aldrich | ab286868 |
| LysoTracker Deep Red | Invitrogen | L12492 |
| VECTASHIELD Antifade Mounting Medium with DAPI | VectorLabs | H-1200-10 |
| **Enzymes** |  |  |
| DpnI (10U/μL) | Thermo Fisher Scientific | ER1701 |
| Phusion High-Fidelity DNA Polymerase (2U/μL) | Thermo Fisher Scientific | F530S |
| **Kits** |  |  |
| GeneJET Plasmid Miniprep Kit | Thermo Fisher Scientific | K0502 |
| Dynabeads™ Protein G | Invitrogen | 10004D |
| **Solutions** |  |  |
| 1X Phosphate-Buffered Saline (PBS) | NaCl: 137mM, KCl: 2.7mM, Na<sub>2</sub>HPO<sub>4</sub>: 10mM, KH<sub>2</sub>PO<sub>4</sub>: 1.8mM – dissolved in 1 litre pH 7.4 |  |
| 1X Tris acetate EDTA (TAE) | 0.4M tris acetate (pH ~8.3), 0.01 MEDTA (Dilute 1:10) using ddH<sub>2</sub>O |  |
| 5X Tris-Glycine-SDS buffer (Gel Running Buffer) | 0.5M Tris, 1.92M Glycine, 0.5% SDS |  |
| 1X Transfer Buffer | Tris (3.03g), Glycine (14.4g), Methanol (200ml), add ddH<sub>2</sub>O to 1L |  |
| RIPA lysis Buffer | 20mM Tris-Cl (pH 7.4), 150mM NaCl, 2% glycerol, 0.5% Triton-X 100, 1.6mM EDTA, supplemented with Protease inhibitor cocktail and 1mM PMSF |  |
| Washing Buffer for co-IP | 20mM Tris-Cl (pH 7.4), 150mM NaCl |  |
| 5X SDS protein loading dye | 250mM Tris-HCl (pH 6.8), 10% SDS, 30% glycerol (v/v), 0.05% Bromophenol Blue (w/v), 500 mM DTT |  |
| 6X DNA loading dye | 30% Glycerol (v/v), 0.25% Bromophenol blue (w/v), 0.25% Xylene cyanol FF (w/v) |  |
| Blocking buffer | 2% Bovine Serum Albumin (BSA) in 1X PBS |  |
| Permeabilization buffer | 0.1% Triton X-100 in 1X PBS |  |
| Fixation buffer | 4% paraformaldehyde in 1X PBS |  |
| Experimental model: Cell lines |  |  |
| HCT116 | NCCS, Pune, India<br>(Cell Repository) | NA |
| THP-1 | NCCS, Pune, India<br>(Cell Repository) | NA |
| RAW 264.7 | NCCS, Pune, India<br>(Cell Repository) | NA |
| HeLa | ATCC | ATCC® CCL-2™ |
| Biological samples |  |  |
| Defibrinated human blood | Voluntary donation<br>(with informed consent<br>through a signed<br>consent form) | NA (In-house collection) |
| Software |  |  |
| Fiji (ImageJ) | NIH | https://imagej.nih.gov/ij/index.html |
| GraphPad Prism 10 | GraphPad Software<br>Inc. | https://www.graphpad.com/ |
| BioRender | BioRender.com | https://www.biorender.com/ |
| Adobe Illustrator | Adobe Inc. | https://adobe.com/products/illustrator |
| DeltaVision SoftWorx software | GE Healthcare | https://www.gehealthcare.com/ |
| Equipment |  |  |
| DeltaVision Elite Deconvolution<br>Microscope | GE Healthcare | NA |
| Varioskan Flash plate reader | Thermo Fisher<br>Scientific | N06354 |
| Gel documentation system | Syngene, USA | G:box |
| Mini Trans-Blot Electrophoretic<br>Transfer Cell | Bio-Rad | 1703930 |
| Countess 3 Automated Cell Counter | Thermo Fisher<br>Scientific | AMQAX2000 |

### Immunofluorescence microscopy

HCT116 and RAW 264.7 cells were seeded on coverslips and allowed to attach overnight. THP-1 cells were seeded on coverslips and treated with 20 ng/mL PMA for 48h to promote attachment. After attachment, cells were infected and/or treated with the indicated compound (Table 5) for 6h. Cells were then fixed using a fixation buffer (4% paraformaldehyde in 1X PBS). For antibody staining, cells were permeabilized with a permeabilization buffer (0.1% Triton X-100 in 1X PBS) for 10 min, followed by blocking with a blocking buffer (2% BSA in 1X PBS) for 1h at room temperature. Cells were incubated with the primary antibodies (listed in Table 4) overnight at 4°C, followed by a 1h incubation with the secondary antibodies (listed in Table 4) at room temperature in the dark. Phalloidin (Sigma-Aldrich, 68825) dye was then added at a 1:60 dilution and incubated for 1h at room temperature in the dark. When combined with immunostaining, phalloidin dye was co-incubated with the secondary antibody under the same conditions. Coverslips were mounted using Vectashield antifade reagent with DAPI (VectorLabs, H-1200-10), and fluorescence was observed using a DeltaVision Elite Deconvolution Microscope.

### Measurement of lysosomal acidity and hydrolytic activity

For fixed-cell microscopy, cells were stained with 100nM LysoTracker Deep Red (Invitrogen, 12492) for 30 min at 37°C in 5% CO₂. After staining, cells were washed with 1X DPBS, fixed with fixation buffer, and imaged. DQ-BSA (Abcam, ab286868) was added at a concentration of 16µg/mL along with 50nM LysoTracker in DMEM supplemented with 0.5% FBS and incubated for 1h. Cells were then washed twice with 1X DPBS, and a live-cell imaging solution (Invitrogen, A59688DJ) was added. Live-cell snapshots were acquired.

### Immunoblotting

HCT116 cells were seeded in 12-well plates and incubated overnight for attachment. Subsequently, cells were either infected for 3 and 6h or treated with rLLO (Abcam, ab83345) for 2h, in the presence or absence of Torin1 (Tocris Bioscience, 4247) or CQ (Sigma-Aldrich, C6628). After the specified treatment durations, cells were washed with ice-cold PBS and lysed in 1X SDS protein loading dye using a cell scraper on ice. The resulting cell lysates were then boiled at 99°C for 15min. Subsequently, the samples were separated by SDS-PAGE and transferred to PVDF membranes (PALL Life Sciences, 79548A) via wet transfer method. The membranes were incubated with primary antibodies (listed in Table 4) overnight at 4°C, followed by a 1h incubation with HRP-conjugated secondary antibodies (listed in Table 4) at room temperature. Protein bands were visualized using an enhanced chemiluminescence substrate (Clarity Bio-Rad, 170-5061), and images were captured using a gel documentation system. Finally, band intensities were quantified using ImageJ software.

### Bacterial growth curve in BHI broth

A single colony was inoculated in BHI broth and incubated overnight at 37°C without shaking. The following day, a secondary culture was prepared by diluting the overnight culture 1:10 in fresh BHI broth. Once the culture reached an OD_600_ of 0.8 to 1.0, a total of 100µl of culture was added to each well of a 96-well plate, ensuring that the final OD_600_ of the added culture was ∼ 0.1 to 0.3 for all bacterial strains. Absorbance at 600 nm was then measured every hour for 12 or 24h using a Varioskan Flash plate reader (Thermo Fisher Scientific).

### Measurement of LLO-mediated hemolytic activity

*Lm* strains were grown in 2.5 mL BHI broth at 37°C overnight without shaking. Stationary-phase cultures were diluted 1:10 in 5 mL fresh BHI broth and incubated at 37°C with shaking until OD_600_ reached 0.8 to 1.0. 1 mL of each culture (normalized to 0.8 to 1.0 OD_600_) was centrifuged at 13,000 x g for 10 min. The human red blood cells (RBCs) (voluntary donation with informed consent through a signed consent form) were washed twice with 1X PBS (pH 7.4) and resuspended to a final concentration of 5% (v/v) in 1X PBS. Supernatants from bacterial cultures were collected and supplemented with 1 mM dithiothreitol (DTT) (Thermo Fischer Scientific, R0862) for 10min at room temperature. In a 96-well plate, 100µL of the bacterial supernatant and 100µL of the 5% RBC suspension were combined (1:1 ratio, 200µL total volume). The mixtures were incubated at 37°C for 45min without shaking. Following incubation, plates were centrifuged, and hemoglobin release in the supernatant was quantified by measuring absorbance at 550nm. Hemolytic activity was determined by quantifying the lysis of RBCs induced by LLO present in culture supernatants.

### CellTiter-Glo (CTG) luminescent cell viability assay

HCT116 cells were counted using the Countess 3 Automated Cell Counter, and equal numbers (10,000 cells per well) were plated on a 96-well plate. A dose-response assay for rLLO (ranging from 2.5µM to 20µM) was performed for 2, 4, and 6h. At each incubation time point, 10µL of CellTiter-Glo reagent (Promega, G7570) was added to each well, and luminescence was measured using the Varioskan Flash plate reader (Thermo Fisher Scientific).

### esiRNA transient transfection

HCT116 were plated in a 24-well plate at a confluency of ∼ 30%. Cells were transfected with 400ng of TFEB esiRNA (Sigma-Aldrich, EHU059261) using Lipofectamine 2000 (Invitrogen, 11668019) transfection reagent for 48h, universal negative control siRNA (Sigma-Aldrich, SIC001) was used as control. After 48h, cells were subjected to intracellular CFU infection assay.

### Trichloroacetic acid (TCA) precipitation

To quantify the secretory levels of LLO from the supernatant of *Lm* cultures grown in BHI broth, TCA (Thermo Fischer Scientific, 76-03-9) precipitation was performed. *Lm* strains were first grown to stationary phase in BHI broth overnight. The cultures were then diluted 10-fold in fresh BHI broth and incubated at 37°C with shaking until they reached an OD600 of 0.8-1.0. For sample preparation, 1 mL of culture was transferred into a microcentrifuge tube and centrifuged at 13,000 x *g* for 15 min. The bacterial pellet was resuspended in 1 X SDS loading dye to prepare samples for intracellular LLO analysis, while the supernatant was mixed with 100µL of 100% TCA and incubated on ice for 1h to precipitate secreted proteins. After incubation, the mixture was centrifuged at 13,000 x *g* for 10 min, and the resulting protein pellet was washed twice with ice-cold acetone. The final pellet was resuspended in 1 X SDS loading dye containing 0.1N NaOH.

### Co-immunoprecipitation

HCT116 cells were cultured in 100 mm dishes. Monolayers were trypsinized, washed twice with 1 X DPBS, and centrifuged at 600 x *g* for 5 min. The resulting cell pellets were lysed in ice-cold RIPA buffer (20 mM Tris-Cl, pH 7.4; 150 mM NaCl; 2% glycerol; 0.5% Triton-X 100; 1.6 mM EDTA) supplemented with a protease inhibitor cocktail (Roche, COEDTAF-RO) and 1 mM PMSF (Sigma-Aldrich, P7626). Lysates were vortexed and sonicated twice for 30 seconds each, with 1 min incubation on ice between sonication steps. Following centrifugation at 12,000 x *g* for 30 min at 4°C, supernatants were collected. Bacterial supernatant was obtained from a secondary culture grown to an OD_600_ of 1. Bacterial cells were removed by centrifugation at 14,000 x *g* for 10 min, and the supernatant was filtered through a 0.22µm pore-size filter for further use. For co-immunoprecipitation, mammalian protein lysate and bacterial supernatant were incubated with anti-LC3 antibodies (MBL Life Sciences, M152-3), following the manufacturer’s recommendations. Protein-antibody complexes were captured using Protein G Dynabeads (Invitrogen, 10004D) and incubated overnight at 4°C with rotation. After incubation, beads were washed three times with washing buffer (20 mM Tris-Cl (pH 7.4), 150 mM NaCl). Finally, the beads were then resuspended in 1X SDS protein loading dye and subsequently analyzed by SDS-PAGE and western blotting.

### Live cell imaging

HeLa cells (ATCC) were cultured and transfected with Lifeact-mCherry (Addgene plasmid #193300) using Lipofectamine 2000 transfection reagent (Invitrogen, 11668019). After 24h of transfection, cells were infected with GFP-tagged WT *Lm* as per the previously described infection protocol. Live cell imaging was carried out at 6hpi using a DeltaVision Elite Deconvolution Microscope at a frame rate of 1 frame every 1s for total duration of 1.3min (eight Z sections of 0.2µm thickness).

### Image analysis

Intracellular motility dynamics of *Lm* were quantified using the TrackMate plugin in ImageJ ^103^. Xenophagosomes (identified by the colocalization of LC3 and *Lm*) and xenolysosomes (identified by the triple colocalization of LC3, LAMP1, and *Lm*) were manually quantified using the Arbitrary line profile tool in Softworx software on individual z-stack images. The same approach was used to analyze the recruitment of p62 to *Lm*, as well as the colocalization of LAMP1 and LysoTracker or DQ-BSA with *Lm*. Actin tail formation and actin cloud association with *Lm* were quantified using the Cell Counter plugin in ImageJ. LAMP1-positive puncta were quantified using the Analyze Particles tool in ImageJ. To assess LC3-LAMP1 and LLO-LC3 colocalization, the Colocalization plugin in ImageJ was utilized, followed by particle analysis on the merged signal to estimate the number of colocalized puncta. The fluorescence intensity of LysoTracker and DQ-BSA was determined by quantifying the Integrated Density (RawIntDen) in ImageJ. Similarly, nuclear translocation of TFEB was quantified by measuring RawIntDen within nuclear ROIs.

### Statistical analysis

All statistical analyses and graphical representations were performed using GraphPad Prism 10.0.0. Data are presented as the mean ± SEM. P values were determined using the specific methods outlined in the respective figure legends. P value less than 0.05 (P < 0.05) was considered statistically significant.

### Antibodies, reagents and other resources

Details of all antibodies, reagents and other resources are provided in Table 4.

## Supporting information

Intracellular and intercellular host actin-mediated movement of Lm

## Acknowledgments

This work was supported by the Life Sciences Research Board-DRDO (LSRB-310/BTB/2017), the Science and Engineering Research Board (SERB/MN/4471), the Indian Council of Medical Research (ICMR/RM/4846) and the intramural funds from JNCASR (JNCASR/MBGU/RM) to RM. We extend our sincere thanks to all members of the Autophagy Laboratory for their valuable support and insightful discussions. We are deeply grateful to Prof. Daniel Portnoy and Andrea Anaya Sanchez for their continual support with the plasmid constructs and *Lm* strains. We also thank Dr. Veena Ammanathan, Dr. Keerti, Dr. Kushagra Bansal and Dr. Kesavardana Sannula for their thoughtful input and guidance. We acknowledge the research facilities provided by JNCASR. AP received support through a doctoral fellowship from JNCASR. SB and KSR are thankful to JNCASR for its continued support. PS acknowledges IISER Mohali for funding, and DB is grateful to JNCASR for her Integrated Ph.D. fellowship. MN is thankful to JNCASR for post-doctoral fellowship. All the figures were made using Adobe Illustrator with changes applied uniformly across all tests and controls unless mentioned otherwise in the methods. Schematic representations were made using BioRender through a paid subscription.

## Author contributions

A.P. and R.M. conceptualized the study. A.P. and R.M. developed the methodology. A.P., S.B., P.S., and D.B. conducted the investigation. K.S.R., S.B., P.S., A.P., and M.N. analyzed the data. A.P. and R.M. drafted, edited and reviewed the manuscript, while rest of the authors reviewed it. R.M. supervised the project and acquired funding.

## Declaration of generative AI and AI-assisted technologies in the writing process

During the preparation of this work the authors used ChatGPT to improve the language and readability. After using this tool, the authors reviewed and edited the content as needed and take full responsibility for the content of the publication.

